# Proteostasis in Podocytes: Role of Cargo Receptor Surf4 and Collagen IV Trafficking

**DOI:** 10.64898/2026.09.30.755739

**Authors:** Andrey V. Cybulsky, Joan Papillon, Christian-Landry Konde Bilogui, Rachel Shih

## Abstract

Protein misfolding in the endoplasmic reticulum (ER) of podocytes/glomerular epithelial cells (GECs) induces ER stress and contributes to the pathogenesis of glomerular disease. Activation of IRE1α and consequently the unfolded protein response (UPR) and autophagy sustains proteostasis. The present study examines the role of Surf4 and collagen IV-α5 (COL4A5) trafficking in podocyte proteostasis. Glomerular Surf4 protein expression increased in IRE1α-replete mice with experimental focal segmental glomerulosclerosis (FSGS) compared to podocyte-IRE1α knockout mice with FSGS and control mice. Glomerular Surf4 gene expression was greater in human FSGS and membranous nephropathy compared to healthy controls. In cultured GECs, basal Surf4 expression was greater in IRE1α-replete (control) compared to IRE1α knockout GECs, although ER stress (tunicamycin treatment) did not increase Surf4 despite activating the UPR. Tunicamycin enhanced autophagy (LC3-II) in an IRE1α-dependent manner, and the amount of Surf4 localized in autophagosomes. In GECs, tunicamycin increased the amount and colocalization of endogenous COL4A5 or ectopically-expressed COL4A5-wild type (WT) and Surf4 in autophagosomes. Compared to COL4A5-WT, the human COL4A5-C1567R mutant (which is misfolded) showed enhanced basal colocalization with Surf4 in autophagosomes; colocalization was similar to COL4A5-WT after tunicamycin treatment. Degradation of COL4A5-C1567R was more rapid compared to COL4A5-WT and was comparable to degradation of COL4A5-WT in the presence of tunicamycin. Thus, during ER stress, subsets of Surf4 and COL4A5 are redirected from the secretory pathway to autophagy. Surf4 may help deliver misfolded ER cargo, such as COL4A5 for autosomal degradation, thereby sustaining proteostasis in podocytes.

## Introduction

Podocytes or glomerular visceral epithelial cells (GECs) are highly-differentiated, long-living epithelial cells in the glomerular capillary wall, and podocyte injury is the hallmark of many glomerular diseases. Podocytes, together with the glomerular basement membrane (GBM), play a key role in maintaining glomerular permselectivity [1,2]. Collagen IV-α3-5 is a large complex protein trimer produced by podocytes that is a crucial GBM component and is essential for normal glomerular permselectivity [1,3,4]. In certain preclinical and human glomerulopathies, podocyte injury features impaired protein folding and protein homeostasis (“proteostasis”) within the endoplasmic reticulum (ER). This damages the glomerulus and drives proteinuria [5–9]. Protein misfolding causes “ER stress” and activates the unfolded protein response (UPR), a compensatory signaling network. The UPR with its linkage to macroautophagy (hereafter referred to as autophagy) is especially critical in maintaining proteostasis in podocytes, given their limited regenerative capacity [5–9].

Newly translated proteins destined for secretion or to the plasma membrane (the “secretory pathway”) attain their proper folding conformations with the help of chaperones in the ER. Folded proteins undergo further post-translational modifications in the Golgi [6,10–12]. The vesicular transport of cellular cargo from the ER to the Golgi is mediated by the coat protein complex II (COPII). Many ER cargo proteins interact with Sec23B, a core COPII component [11,13,14]; other proteins may instead employ cargo receptors, e.g. Surf4 (surfeit locus protein 4) or LMAN1 (lectin mannose-binding 1) [14–16]. In the early secretory pathway in the ER, Surf4 selectively recognizes and packages specific cargo proteins into transport vesicles.

The UPR is an adaptive mechanism that maintains ER function, enhances protein folding and alleviates stress caused by misfolded proteins. There are three UPR transducers – activating transcription factor 6, PKR-like ER kinase and inositol-requiring enzyme 1α (IRE1α). Our focus has been on IRE1α, a protein kinase and RNase, and the most evolutionarily conserved UPR transducer [12]. Classically, IRE1α RNase splices X-box binding protein 1 (Xbp1) mRNA to generate a potent transcriptional activator of ER chaperone genes and genes involved in ER-associated degradation of misfolded proteins [12].

There is emerging evidence that autophagy is linked with IRE1α and the UPR [12]. In autophagy, damaged proteins/organelles are engulfed in a double-membraned vesicle (autophagosome), which fuses with a lysosome (autolysosome), and the contents of the vesicle are then degraded by lysosomal hydrolases. A key protein involved in autophagy is microtubule-associated protein 1 light chain 3 (LC3-I), which becomes lipidated (i.e. LC3-II) upon localization to autophagosomes. IRE1α can stimulate autophagy through transcriptional and post-translational mechanisms [12,17]. During ER stress, IRE1α can drive transcription of the autophagy mediators LC3, ATG5, and ATG7 [18]. Proteins encoded by these genes are involved in formation of autophagosomes, which is considered the rate-limiting step in the autophagic process [19].

A stress-induced autophagy pathway called reticulophagy (or ERphagy) ensures optimal ER function by timely removal of damaged ER and its contents from the cell [20–22]. Reticulophagy may also regulate basal ER turnover and is upregulated during nutrient depletion or ER stress [23]. In particular, reticulophagy may involve redirection of COPII vesicles in the secretory pathway towards autophagy. Reticulophagy is mediated by ER resident reticulophagy adaptors, which contain motifs for recognition of LC3 in autophagosome membranes, and deliver ER fragments to autophagosomes. For example, family with sequence similarity 134 member B (FAM134B) and the long isoform of reticulon-3 (RTN3L) are reticulophagy adaptors with reticulon homology domains, which allow them to modulate membrane curvature and facilitate the budding of ER fragments [24]. ER-derived vesicles of the COPII system can assist in fragmentation of the ER and autophagosome growth, and deliver aggregation-prone proteins to autophagosomes [20,21,24,25].

We demonstrated a cytoprotective role for the IRE1α pathway and reticulophagy in glomerular disease [13]. Podocyte-specific knockout (KO) of IRE1α in mice resulted in proteinuria, podocyte depletion, and ultrastructural podocyte abnormalities as mice aged. Deletion of IRE1α exacerbated podocyte injury and proteinuria in experimental focal segmental glomerulosclerosis (FSGS; adriamycin nephrosis), anti-GBM glomerulonephritis (GN) and diabetic nephropathy [18,26,27], and deletion impaired activation of the UPR and autophagy, a critical protective mechanism to clear misfolded proteins in podocytes [6,12,17,22,28–30]. Importantly, our analyses of human glomerular gene expression support activation of the IRE1α-UPR-autophagy axis in humans with GNs [18,31]. Moreover, ER stress markers are increased in glomeruli of kidney biopsies from patients with hereditary GN, acquired GN (FSGS and membranous nephropathy) and diabetic nephropathy [6,32,33]. Thus, in response to injury, activation of IRE1α/UPR preserves glomerular integrity in various preclinical and most likely human glomerulopathies.

In a proteomic analysis using cultured GECs, ER stress induced upregulation of novel targets of IRE1α-Xbp1, including mediators of reticulophagy and cargo receptors (such as Surf4) [13]. Activated IRE1α upregulated Sec23B and RTN3L together with LC3-II in GECs and in experimental FSGS [13]. Remarkably, Sec23B and RTN3L mRNA are also upregulated in human FSGS [13]. We concluded that IRE1α and ER stress can redirect a subset of Sec23B-positive vesicles to autophagosomes, to deliver ER fragments coated with RTN3L [13], in keeping with reticulophagy pathways described earlier [20,21].

The pathway for secretion of GBM collagen IV-α3-5 remains to be fully elucidated, and while the COPII system has been implicated in collagen secretion, there has been some controversy [34]. Recently, it was reported that certain collagens may be cargoes of Surf4 [16]. Complex proteins, such as collagen IV, may misfold basally or during ER stress [10,11]. Moreover, in podocytes, some collagen IV missense mutant proteins may be retained in the ER, induce ER stress and cause human disease (Alport nephritis) [35–38]. We showed that during ER stress in GECs, a portion of collagen IV-α5 (COL4A5) colocalizes with RTN3L and LC3-II (in autophagosomes) in an IRE1α-dependent manner [13], indicating that misfolded COL4A5 may be degraded via reticulophagy, thereby limiting accumulation in podocytes and cytotoxicity.

In the present study, we show that during ER stress, subsets of Surf4 and COL4A5 colocalize in autophagosomes. The human COL4A5-C1567R mutant (which is misfolded) [39] showed enhanced basal colocalization with Surf4 in autophagosomes. Thus, during ER stress, Surf4 may help deliver misfolded ER cargo, such as COL4A5 for autophagosomal degradation.

## Methods

### Antibodies

Rabbit anti-LC3B (2775) mouse anti-LC3B (83506) and rabbit monoclonal anti-HA (C29F4, 3724) antibodies were purchased from Cell Signaling Technology (Danvers, MA). Rat anti-GRP94 (sc-32249) and mouse anti-HA, F7 (sc-7392) antibodies were from Santa Cruz Biotechnology (Santa Cruz, CA). Rabbit anti-mesencephalic astrocyte-derived neurotrophic factor (MANF; PAB13301) was from Abnova (Walnut, CA). Rat anti-collagen IV-α5 (COL4A5) clone H53 (7078) was purchased from Chondrex Inc. (Woodinville, WA). Mouse anti-LC3B clone 5F10 (ALX-830-080-C100) and rabbit anti-calnexin antibody (ADI-SPA-860) were from Enzo Life Sciences (Ann Arbor, MI). Rabbit anti-Surf4 antibody (11599-1-AP) was from Proteintech (Rosemont, IL). Rabbit anti-RTN3L (PA2256) was from Boster Biological Technology (Pleasanton, CA). Rabbit anti-actin (A2066) was from MilliporeSigma (Mississauga, ON). Non-immune IgG and secondary antibodies were from Jackson ImmunoResearch Laboratories (West Grove, PA) or ThermoFisher Scientific.

### Studies in mice

Generation, genotyping and characterization of podocyte-specific IRE1α KO mice was published previously [18,26]. Production of adriamycin nephrosis was described previously [18]. The animal protocol was approved by the McGill University Animal Care Committee, and our studies comply with the guidelines established by the Canadian Council on Animal Care.

### Studies in cell culture

Primary-immortalized non-clonal control (IRE1α-replete) and IRE1α KO GECs were generated and characterized as described previously [18]. In KO GECs, there is an absence of IRE1α mRNA and protein. GECs were cultured in K1 medium or in DMEM/F12 with 5% fetal bovine serum [27]. For experiments, GECs were allowed to proliferate for 1-3 days at 33°C, and were then switched to 37°C to differentiate for 24-48 h. HEK 293T cells were cultured in DMEM with 10% fetal bovine serum. 293T cells were transiently transfected using Lipofecatmine 2000 according to the manufacturer’s instructions.

COL4A5 plasmids (lentiviral expression vectors) were kindly provided by Drs. Jeffrey Miner and Kohei Omachi (Washington University, St. Louis, MO). These included pLV BSD-human HA-COL4A5-wild type (WT), pLV BSD-human HA-COL4A5-C1567R and pLV BSD-human HA-COL4A5-G869R [39]. To transduce control and IRE1α KO GECs, lentivirus stocks were prepared in 293T cells. Briefly, 293T cells were cultured and transfected with plasmids as above. Medium was changed after 24 h. Viruses were collected 48-72 h after the start of transfections. Media containing the COL4A5 lentiviruses were added to GECs 24 h after plating in the presence of polybrene (8 μg/ml), and cells were incubated for 48 h at 33°C. Then, clones were selected by culturing after addition of blasticidin (5 μg/ml) to the medium. Clones were expanded and lysates were tested for COL4A5 expression by immunoblotting.

### Microscopy

For immunofluorescence microscopy, kidney poles were snap-frozen in isopentane (−80°C). Cryostat sections (4 μm thickness) were cut and mounted onto coverslips. GECs were plated on coverslips at 1×10^4^ cells/cm^2^. Tissue sections or cells were fixed/permeabilized with 100% methanol, blocked in 5% bovine serum albumin, and incubated overnight at 4°C with primary antibodies. In control incubations (performed in parallel), primary antibody was omitted or replaced with nonimmune IgG. Next, coverslips were incubated with fluorophore-conjugated secondary antibodies at 22°C for 2 h. In some experiments, nuclei were stained with 4′,6-diamidino-2-phenylindole (DAPI) or Hoechst H33342 (ThermoFisher) for 20 min [13,18,27]. Glomeruli or cells in experimental groups were selected randomly and images were acquired using a Zeiss Axio Observer fluorescence microscope with visual output connected to an AxioCam MRm monochrome camera. To allow comparisons of fluorescence intensity, images were taken at the same exposure time. Final images were collected from series of images derived from different focal planes (Z-stack) and subjected to deconvolution, extended depth of focus and maximum projection using Zen 2.6 Pro Zeiss software. Quantification of fluorescence intensity was done with the histogram function of ImageJ software (National Institutes of Health, Bethesda, MD), and results are expressed in arbitrary units [13,18,27]. The glomerular fluorescence intensity was normalized to the total fluorescence in each image.

The ImageJ protocol to measure puncta formation and particle size was carried out as previously [13]. ImageJ allows definition of autophagosome particle size and the threshold of immunofluorescence intensity to count autophagosomes and estimate cell size. Color photographs were split into 3 separate channels and converted to grayscale. Cell contours were selected using the freehand tool on grayscale images, and the area of the cross-section was measured. Image histogram plots were obtained, representing the range of pixel brightness values in each image. Threshold intensity was set at the end of the histogram declining slope (thereby only including the brightest fluorescent signals and omitting cellular features and background). Previous studies of autophagosomes demonstrated that most particles are between to 0.5-25 μm^2^. The ImageJ plugin JACoP was used to evaluate colocalization in the thresholded images [13,40]. We present the Manders’ overlap coefficient (MOC) [13].

### Immunoblotting

The protocol for immunoblotting was described earlier [26]. Chemiluminescence was detected in a ChemiDoc Touch Imaging System (Bio-Rad; Mississauga, ON); signal saturation was monitored with Image Lab (Bio-Rad) and only signal intensities within a linear range were analyzed. Band densitometries were quantified using ImageJ and values were normalized to the expression of β-actin.

### Gene expression analysis

The publicly accessible dataset GSE108109 was used for the expression analysis of glomerular genes [31,41]. The dataset contains microarray gene expression data (mRNAs) of microdissected whole glomeruli from human kidney biopsies. The data are presented as the fold-increase of gene expression in disease above healthy control. The P-values, which we present in this manuscript, are those calculated and reported within the datasets by their original authors.

### Statistical analysis

Values are expressed as mean ± standard error. Data were processed in Prism (GraphPad Software, La Jolla, CA). In experiments with three or more groups, one way analysis of variance (ANOVA) was used to determine significant differences among groups; where relevant, additional comparisons were calculated and P values were adjusted according to the Holm-Sidak method. Significant differences between two groups are displayed with lines between columns, with asterisks denoting P values. In the absence of such lines, differences were not statistically significant.

## Results

### Glomerular Surf4 expression is enhanced in glomerulopathies

In a previous study involving discovery proteomics in GECs, we observed that expression of Surf4, an ER-COPII vesicle cargo receptor, was increased during TM-induced ER stress in an IRE1α-dependent manner [13]. We also showed that during ER stress, the secretory pathway can be redirected to reticulophagy [13], and thus, the present study examined whether Surf4 may be involved in autophagy. First, we examined glomerular expression of Surf4 in experimental and human glomerulonephritis, which features podocyte injury. In mice with adriamycin nephrosis (experimental FSGS), glomerular Surf4 protein expression was increased significantly in control mice (IRE1α-replete) compared to podocyte-specific IRE1α KO mice (Fig. 1A-B). This increase was evident at 4 weeks, a time point at which control mice showed 20-fold greater albuminuria compared to IRE1α KO mice [18]. Glomerular Surf4 gene expression was increased in two human glomerulopathies that involve podocyte injury, i.e. FSGS and membranous nephropathy compared to healthy controls (Fig. 1C). The Human Protein Atlas reports that basal Surf 4 protein expression is enriched in the normal human glomerulus compared to renal tubules [42].

**Figure 1.**
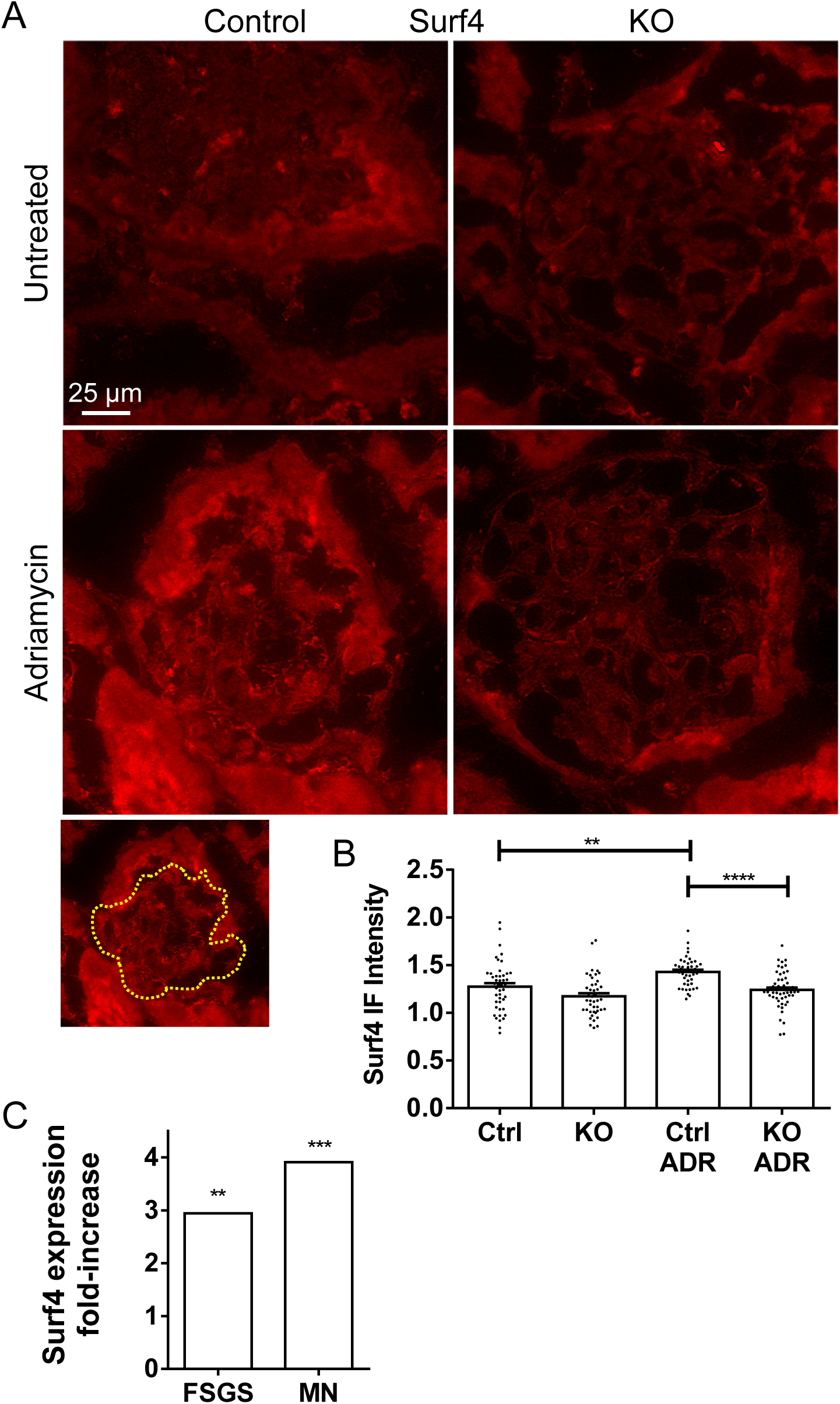
Glomerular Surf4 expression is enhanced in glomerulopathies. A and B) Glomerular Surf4 protein expression is increased significantly in control (Ctrl) mice with adriamycin (ADR) nephrosis compared to podocyte-specific IRE1α KO mice. Immunofluorescence (IF) staining with anti-Surf4 antibody was performed at 4 weeks. An additional adriamycin photo (reduced size) outlines the glomerulus, which had been localized by staining with anti-synaptopodin antibody (not shown). 4 mice per group. N≥45 glomeruli per group. C) Glomerular Surf4 gene expression is increased in human FSGS and human membranous nephropathy (MN) compared to healthy controls. Dataset GSE108109. Results are presented as fold-increase above control. FSGS N=30, MN N=44, control N=6. **P<0.01, ***P<0.001, ****P<0.0001.

### IRE1α regulates basal Surf4 expression in GECs

We induced ER stress by incubating control (IRE1α-replete) and IRE1α KO GECs with tunicamycin (TM), for 6 or 24 h. TM reduces N-linked glycosylation and causes protein misfolding. Basal Surf4 expression was greater in control compared to IRE1α KO GECs (Fig. 2). TM did not increase Surf4 expression above basal levels, but increased the ER chaperones GRP94 and MANF at 24 h, confirming activation of the UPR. MANF (but not GRP94) induction was reduced significantly in IRE1α KO GECs, and in control GECs treated with 4μ8C, an inhibitor of the IRE1α RNase [18]. Thus, GRP94 induction may be more dependent on ATF6, while MANF is dependent on IRE1α.

**Figure 2.**
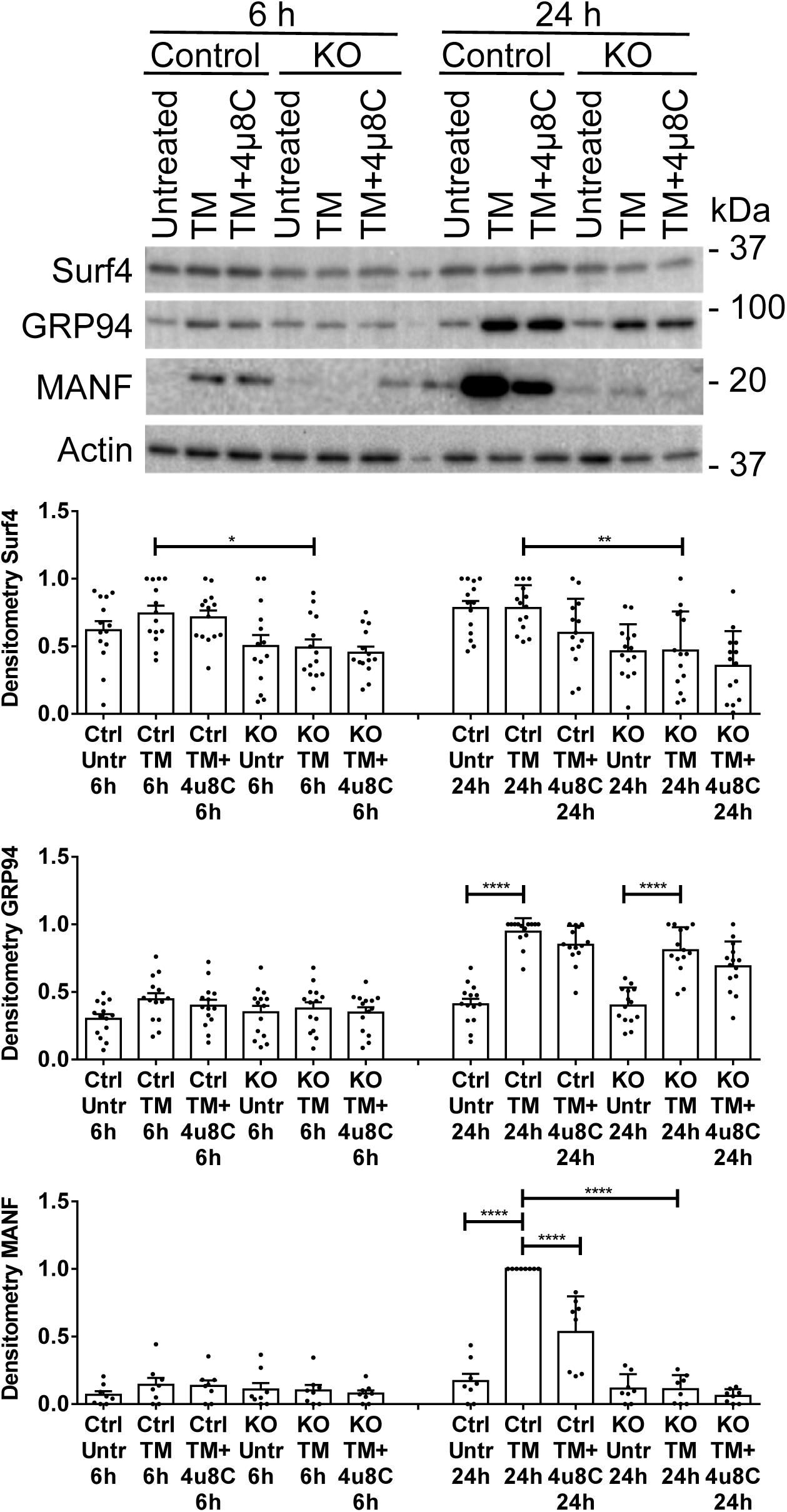
IRE1α regulates Surf4 expression. ER stress was induced by incubation of control and IRE1α KO GECs with TM (1μg/ml), for 6 or 24 h. Lysates were immunoblotted with antibodies to Surf4, GRP94 or MANF (representative immunoblots). Signals were quantified by densitometry. Basal Surf4 expression was greater in control compared to IRE1α KO GECs, but Surf4 expression was not increased above basal levels by TM. TM increased expression of the ER chaperones GRP94 and MANF at 24 h. MANF induction was reduced significantly in IRE1α KO GECs, and in control GECs treated with 4μ8C (0.8μg/ml), an inhibitor of the IRE1α RNase. Untr, untreated. SurF4 and GRP94 – 7 experiments; MANF – 4 experiments. *P<0.05, **P<0.01, ****P<0.0001.

### Tunicamycin increases LC3-II and Surf4 in autophagosomes

Control and IRE1α KO GECs were incubated with TM to induce ER stress. Chloroquine was added to block degradation of autophagosomes, enabling the monitoring of autophagic flux. GECs were stained with antibodies to LC3 and Surf4, and particles consistent with the size of autophagosomes were measured (see Methods). In the context of autophagosomes, LC3 staining reflects LC3-II (LC3-I is distributed throughout the cytosol) [13]. In keeping with previous results, TM increased LC3-II particle areas in control and IRE1α KO GECs (Fig. 3 and Suppl Fig. 1A). The magnitude of the increases in LC3-II particle areas were smaller in KO GECs compared to control, indicating a partial dependence on IRE1α (Suppl Fig. 1B). In the presence of chloroquine, there was an increase in basal LC3-II, and TM further increased LC3-II particle areas, although the TM-induced changes were not as large as those without chloroquine (Fig. 3 and Suppl Fig. 1B). Again, the magnitude of these increases in LC3-II particle areas were smaller in KO GECs compared to control. In Suppl Fig. 1C, we also present changes in the number of puncta. Generally, these paralleled the changes in puncta area and were greater in control compared to KO GECs. However, sometimes the area of individual puncta increased without a major increase in the number of puncta, and going forward, we therefore present puncta area. Finally, the increases in LC3-II puncta areas induced by chloroquine and differences between control and IRE1α KO GECs support the view that the puncta indeed represent autophagosomes.

**Figure 3.**
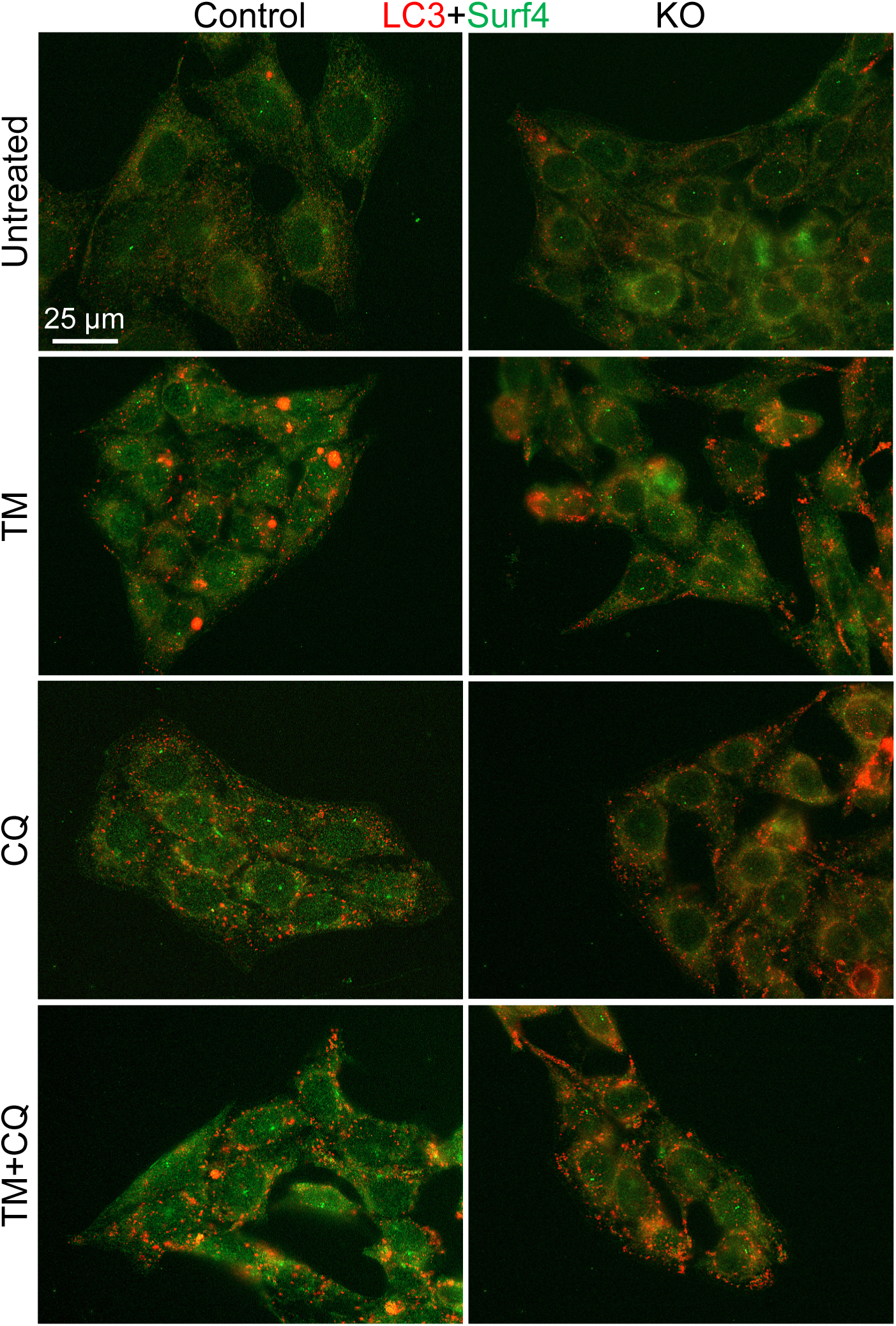
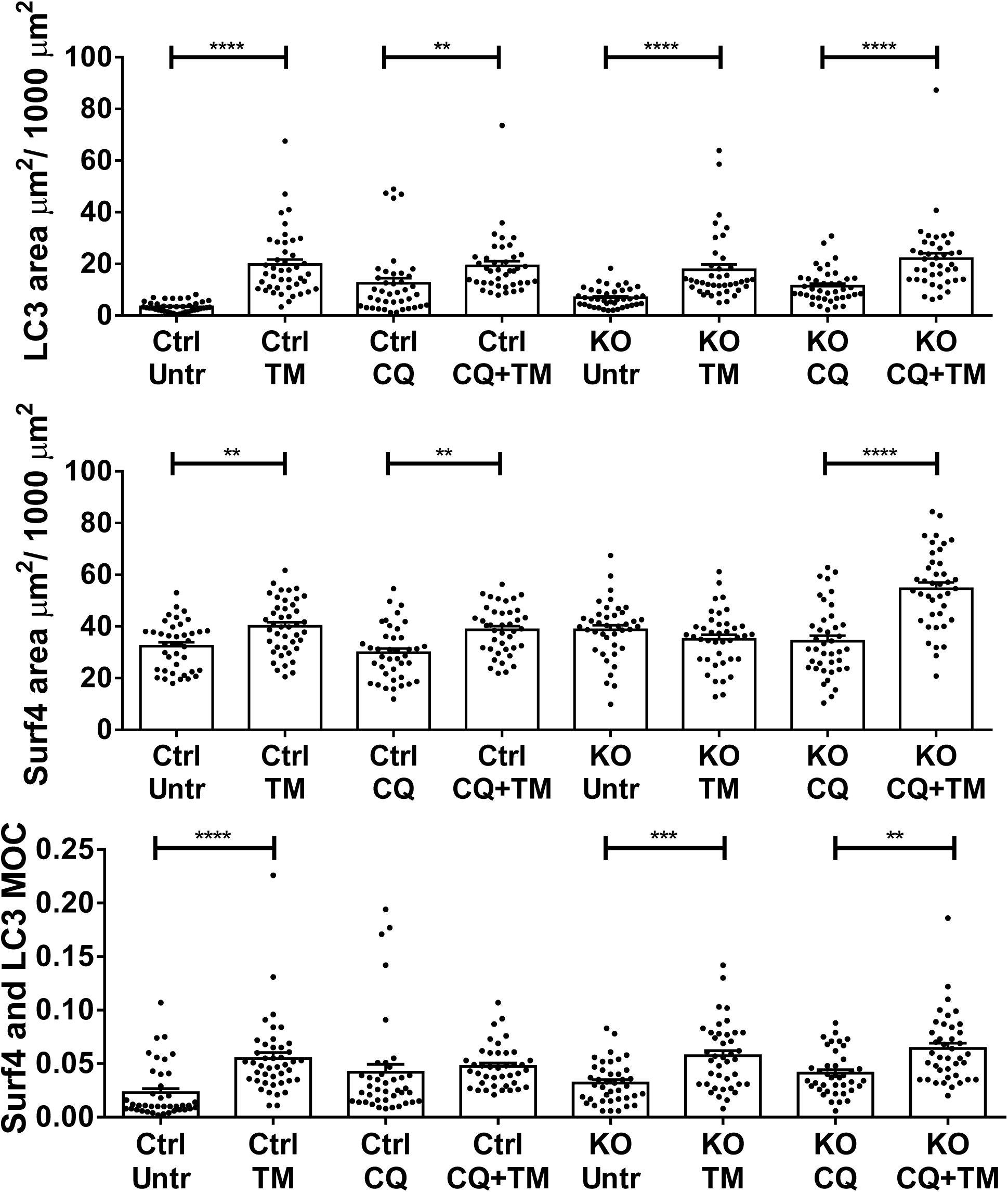
TM increases LC3-II (red) and Surf4 (green) particle areas, as well as colocalization of LC3-II and Surf4. Control and IRE1α KO GECs were untreated, or were incubated with TM (0.25 μg/ml) in the presence or absence of chloroquine (CQ; 25 μM, 24 h). GECs were stained with antibodies to LC3 and Surf4 (representative photomicrographs). Particle areas and colocalization were measured (see Methods). TM increased LC3-II and Surf4 particle areas in control GECs, and less consistently in IRE1α KO GECs. Furthermore, the increases in LC3-II particle area were significantly smaller in IRE1α KO GECs compared to control GECs (see Suppl Fig. 1). CQ, chloroquine. 3 experiments, N=40-43 per group. **P<0.01, ***P<0.001, ****P<0.0001.

TM increased Surf4 particle areas in control GECs, and less consistently in IRE1α KO GECs; i.e. increase was seen only in the presence of chloroquine (Fig. 3 and Suppl Fig. 1A). With TM, there were also increases in colocalization of LC3-II and Surf4 (although this did not reach statistical significance in control GEC in the presence of chloroquine). Together, the results support the view that during ER stress, a subset of Surf4 is redirected to autophagosomes.

### Tunicamycin increases Surf4 and COL4A5 in autophagosomes

In our previous study, after GECs were treated with TM, a portion of COL4A5 colocalized with LC3-II (in autophagosomes, reflecting reticulophagy) [13]. Most likely, this portion of COL4A5 protein reflects misfolded COL4A5. Surf4 is a component of the ER, and a recent study proposed that certain collagens may be cargoes of Surf4 [16]. Thus, we evaluated if during ER stress, COL4A5 might colocalize with Surf4 in autophagosomes (via reticulophagy). We observed that TM increased COL4A5 and Surf4 particle areas, and the colocalization of Surf4 with COL4A5 in control and in IRE1α KO GECs (Fig. 4 and Suppl Fig. 2). Thus, during ER stress, a portion of the COL4A5 and Surf4 localize in autophagosomes. Although IRE1α regulated basal Surf4 expression in GECs (Fig. 2), in these experiments, there was no significant differences in Surf4 and COL4A5 in autophagosomes between control and IRE1α KO GECs.

**Figure 4.**
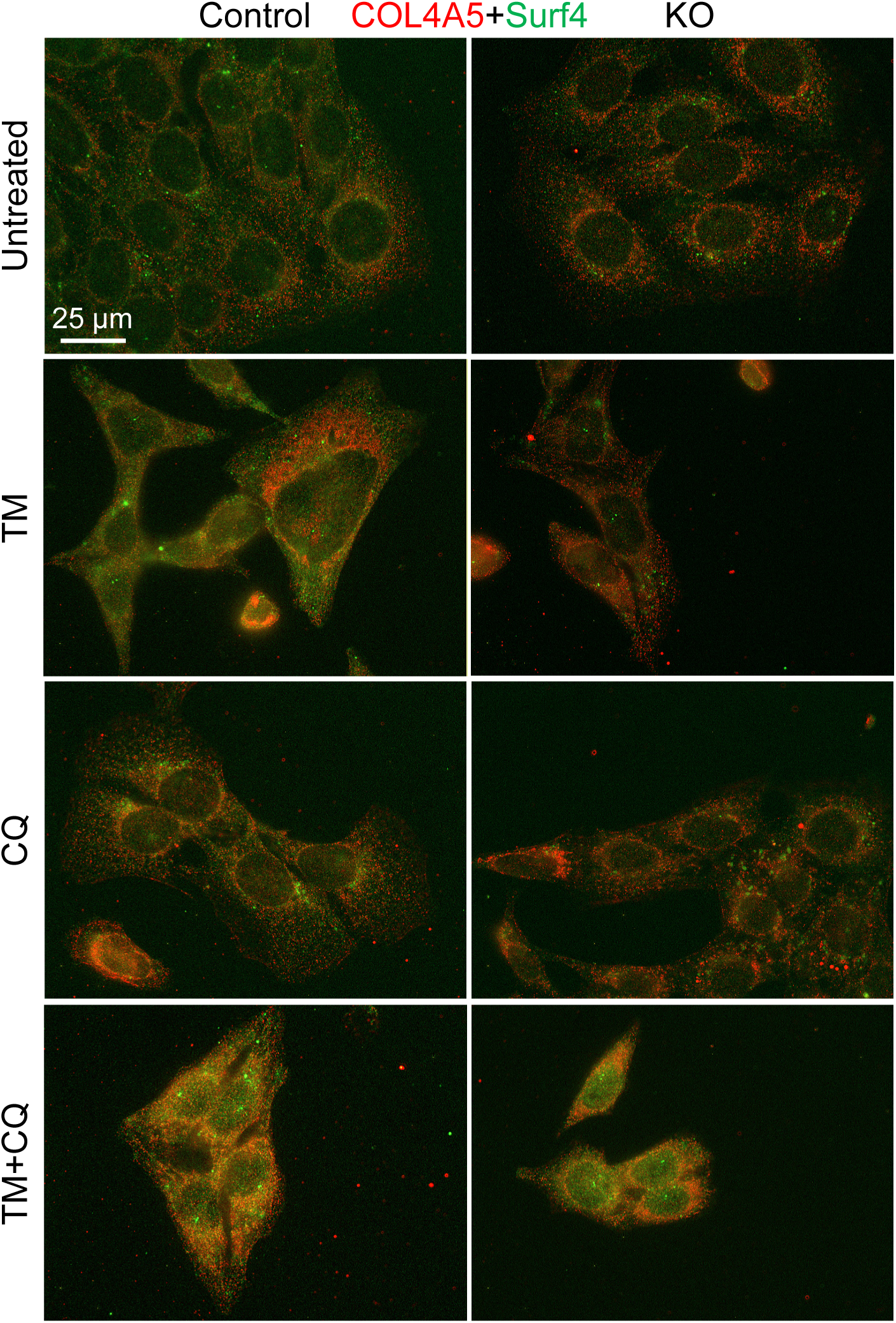
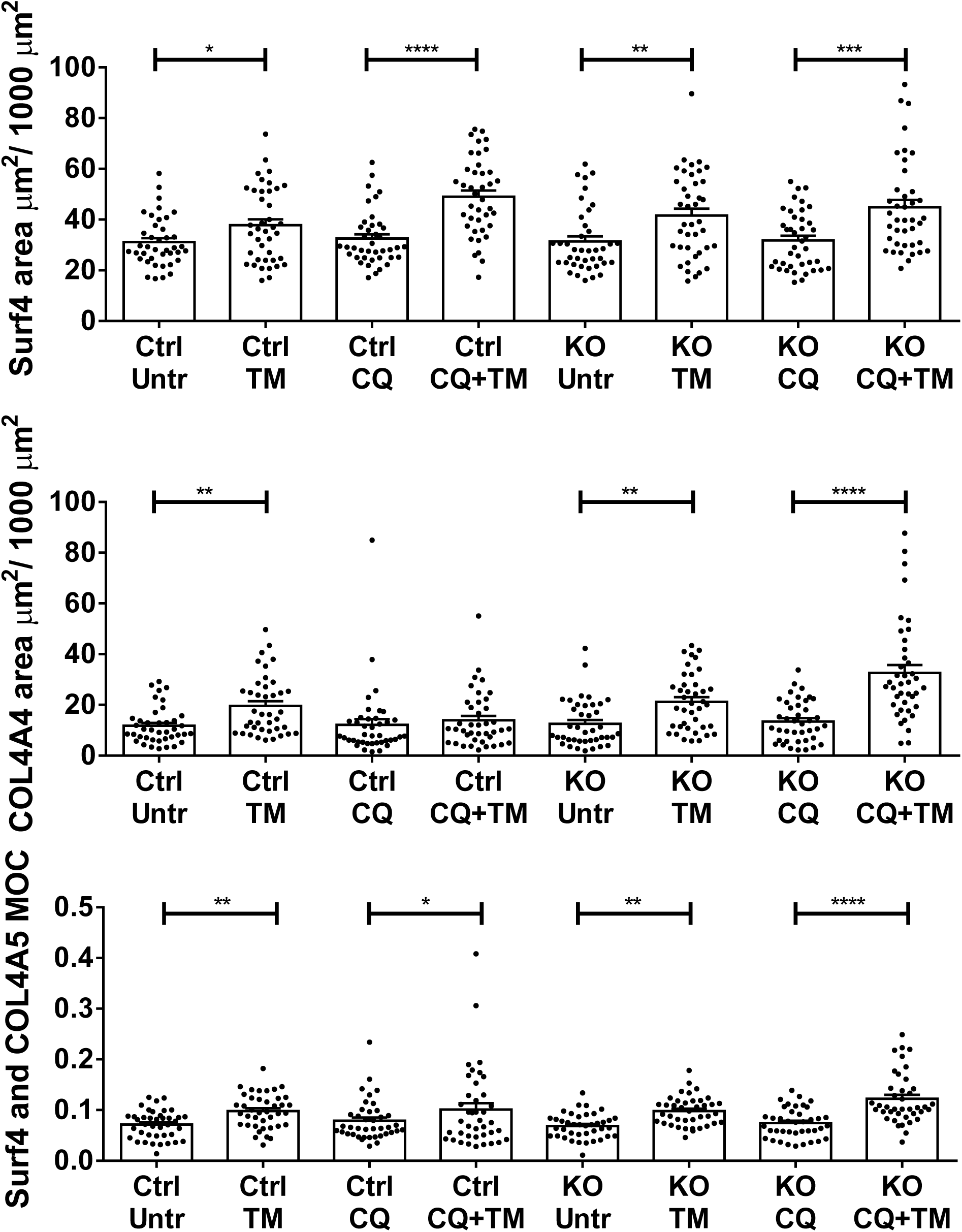
TM increases COL4A5 (red) and Surf4 (green) particle areas, and colocalization of Surf4 with COL4A5 in control and IRE1α KO GECs. Incubations were as in Fig. 3. GECs were stained with antibodies to COL4A5 and Surf4 (representative photomicrographs). 2 experiments, N=38-41 per group.*P<0.05, **P<0.01, ***P<0.001, ****P<0.0001.

To determine if COL4A5 could be associated with Surf4, we investigated potential interactions of the COL4A5 non-collagenous-1 (NC1) domain with Surf4 using Alphafold [43]. (The collagenous domain is too large and disordered for this type of analysis.) The Alphafold structural model predicted NC1 trimer folding and assembly with high confidence, Surf4 folding with moderate to high confidence, and a plausible interaction between Surf4 and the COL4A5 NC1 trimer; however, the Surf4 docking orientation and binding site(s) were uncertain and predicted with low confidence (Suppl Fig. 3). Nevertheless, the model suggests that Surf4 may act as a receptor/chaperone for NC1 because a NC1 trimer forms normally in the presence of Surf4, Surf4 associates with the assembled NC1 domain rather than disrupting it, and the interface spans all three NC1 monomers, which could be biologically important for recognition of a trimeric assembly.

### COL4A5 is degraded by autophagy during ER stress

To provide functional evidence for autophagy degrading COL4A5 during ER stress, control GECs were incubated with cycloheximide, which blocks protein synthesis. GECs were then treated with TM with or without chloroquine (to inhibit autophagy), or for comparison MG132 (to inhibit proteasomal degradation). Results were compared to chloroquine or MG132, without TM. TM+cycloheximide decreased COL4A5 immunofluorescence staining intensity significantly compared to cycloheximide alone (Fig. 5), in keeping with the result of an earlier study, which showed an analogous decrease using immunoblotting [13]. Furthermore, the TM-induced decrease in COL4A5 was attenuated significantly by chloroquine, but not MG132 (Fig. 5). The results are in keeping with a role for autophagy in degradation of COL4A5.

**Figure 5.**
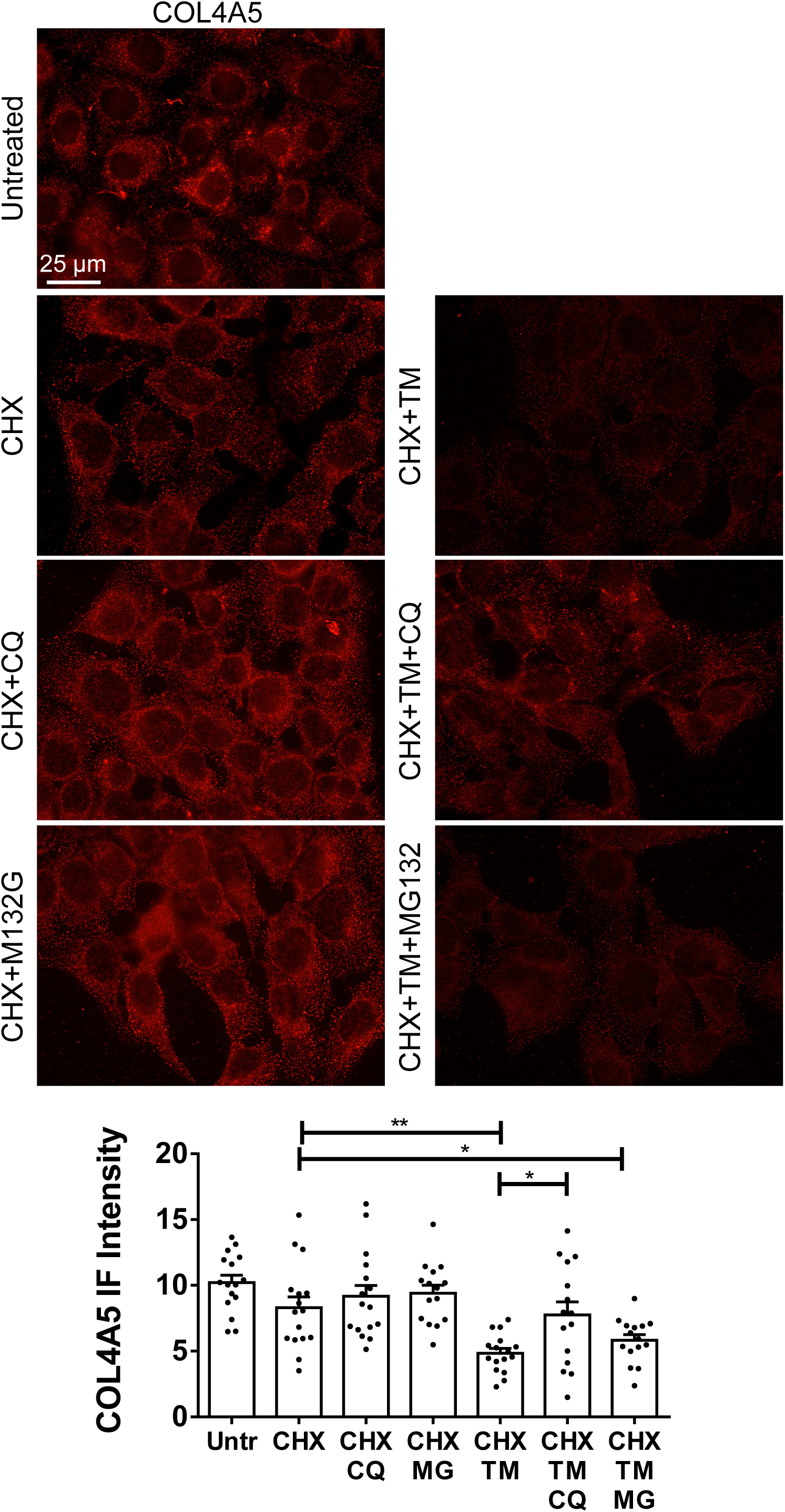
COL4A5 is degraded by autophagy during ER stress (control GECs). GECs were untreated or incubated with TM (1 μg/mL), chloroquine (CQ) (25 μM), cycloheximide (CHX) (1 μM) and the proteasomal inhibitor (MG132) (400 nM, 24 h). GECs were then immunostained with anti-COL4A5 antibody. Immunofluorescence intensity was quantified using the histogram function of ImageJ software. Representative staining and quantification. 3 experiments with 5 photos per group per experiment. * P<0.05, ** P<0.01.

### Human COL4A5 mutants appear in autophagosomes

To substantiate that TM-induced misfolded COL4A5 is transported to autophagosomes, further studies were carried out with COL4A5 mutants that misfold, i.e. cannot assemble into collagen IV trimers [39]. Two COL4A5 mutants, C1567 and G869R, despite misfolding, were expressed comparably to COL4A5-WT when transiently transfected in HEK 293T cells (Suppl. Fig. 4A-B). Furthermore, COL4A5-WT and G869R were secreted from cells into the culture medium, but C1567R was not (Suppl Fig. 4A) and [39]. Transfected 293T cells were untreated or treated with TM and stained with antibodies to HA (to identify epitope-tagged COL4A5) and LC3. Colocalization of HA and LC3-II in puncta was observed in WT-expressing cells treated with TM and in cells expressing the mutants (Suppl Fig. 4C), implying colocalization in autophagosomes. Thus, although COL4A5 mutations impair collagen IV trimerization and the COL4A5 C1567R mutant is not secreted from cells [39], the mutant proteins appear in autophagosomes.

We then transduced control (Suppl Fig. 4D) and IRE1α KO GECs (not shown) with lentiviruses to generate clones stably expressing HA-COL4A5-WT and HA-COL4A5-C1567R. We were unable to propagate GECs stably expressing the G869R mutant beyond two passages. In the experiments presented above (Figs. 3 and 4), the TM-induced changes in puncta were robust even in the absence of chloroquine; therefore, further experiments were performed without adding chloroquine to cells. In control GECs, COL4A5-C1567R expression showed enhanced basal COL4A5, Surf4 and LC3-II puncta area, and enhanced colocalization with Surf4 and with LC3-II compared to COL4A5-WT (Fig. 6 and Suppl Figs. 5 and 6). TM increased COL4A5, Surf4 and LC3-II puncta in HA-COL4A5-WT cells, similarly to the effect of TM on endogenous COL4A5 (Fig. 3) and [13]. Puncta area and colocalization of COL4A5-C1567R with Surf4 and with LC3-II were similar to COL4A5-WT treated with TM (Fig. 6).

**Figure 6.**
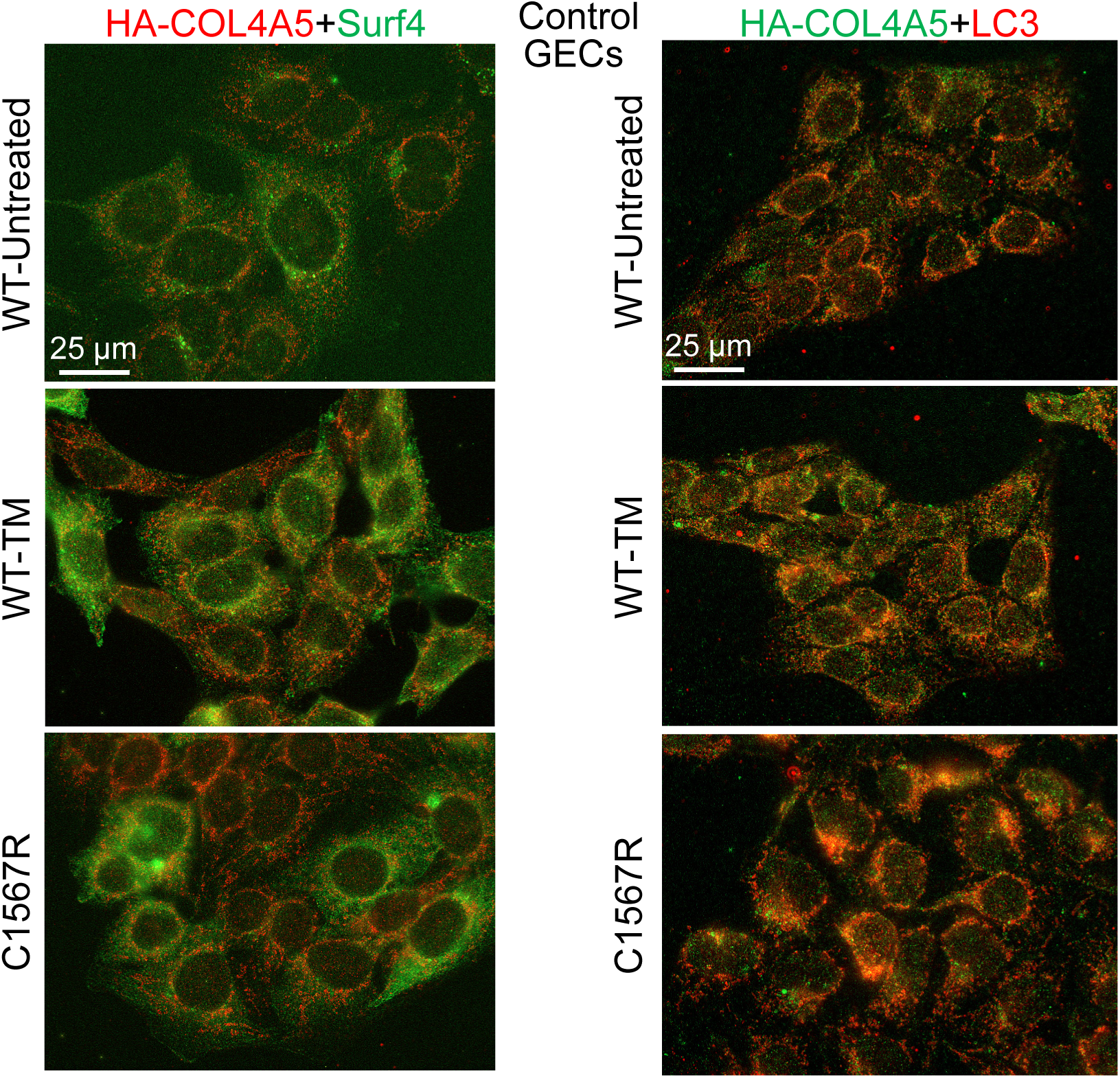
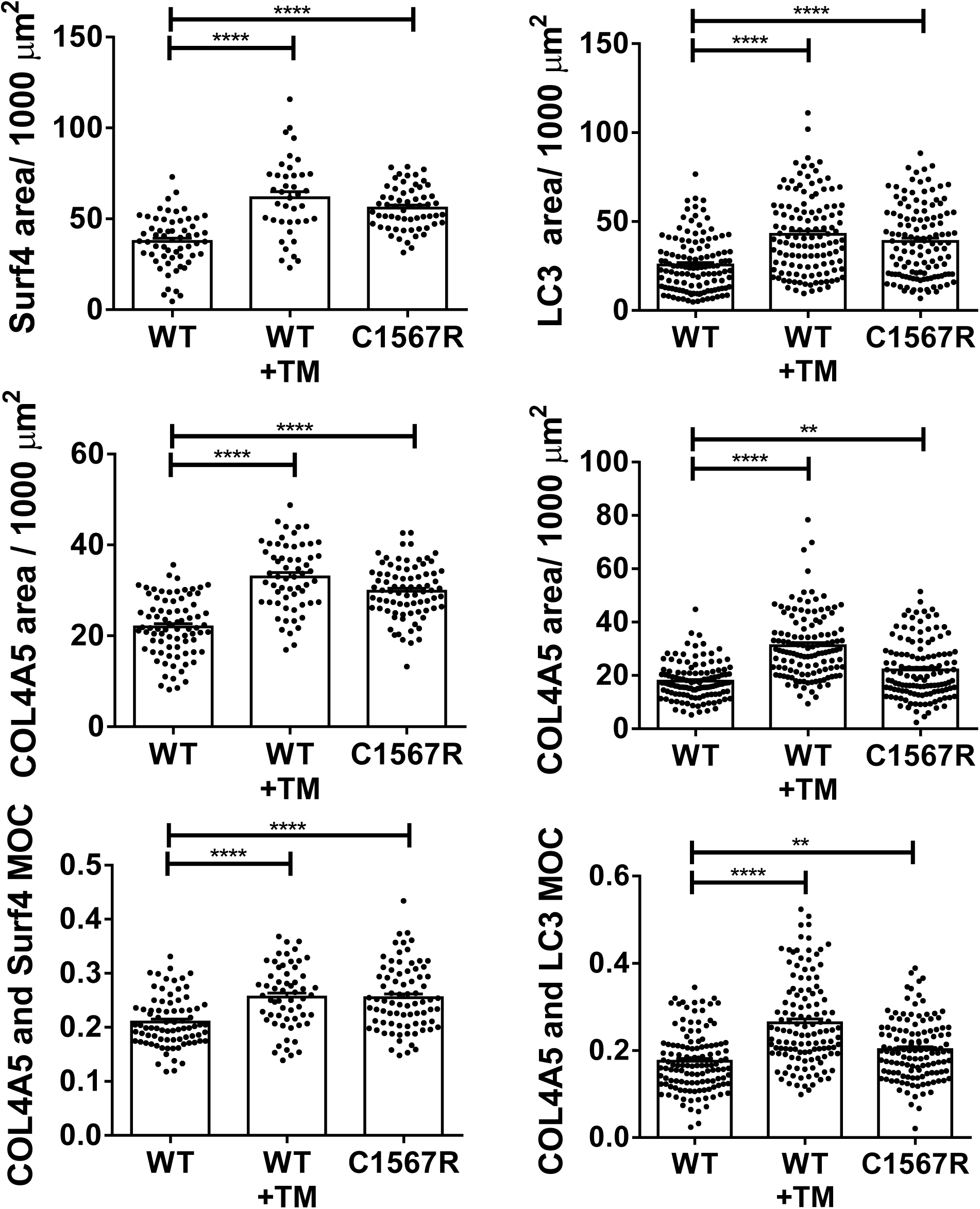
Human COL4A5-C1567R mutant (which is misfolded) shows enhanced basal puncta area and colocalization with Surf4 and with LC3-II compared to COL4A5-WT (control GECs). Puncta area and colocalization of COL4A5-C1567R are similar to COL4A5-WT treated with TM. Control GECs that express HA-COL4A5-WT or HA-COL4A5-C1567R mutant were untreated, or were incubated with TM (0.25 μg/ml; 24 h). GECs were stained with antibodies to HA (COL4A5; red) and Surf4 (green; left) or HA (COL4A5; green) and LC3-II (red; right; representative photomicrographs). COL4A5 and Surf4 (left): 4 experiments, N=61-81 per group. COL4A5 and LC3-II (right): 5 experiments, N=125 per group. **P<0.01, ****P<0.0001.

In IRE1α KO GECs, COL4A5-C1567R showed basal puncta area (COL4A5, Surf4 and LC3-II), comparable to COL4A5-WT in untreated KO cells (Suppl Figure 7). As this result is distinct from the control GECs (Fig. 6), it suggests that IRE1α activity facilitates autophagy.

### COL4A5-C1567R mutant colocalizes with RTN3L

We had demonstrated that during ER stress in GECs, endogenous COL4A5 appeared in autophagosomes together with the reticulophagy adaptor RTN3L [13]. In the present study, we observed that TM increased colocalization of ectopic HA-COL4A5-WT with RTN3L in autophagosomes compared to HA-COL4A5-WT in untreated cells (Suppl Figure 8). Colocalization of COL4A5-C1567R with RTN3L was greater than COL4A5-WT and was similar to COL4A5-WT treated with TM (Suppl Figure 8). Puncta area of COL4A5-C1567R tended to increase above COL4A5-WT, but did not reach statistical significance. The results are consistent with the view that RTN3L is a reticulophagy adaptor for both COL4A5-WT and COL4A5-C1567R.

### COL4A5-C1567R mutant shows accelerated degradation

Since the COL4A5-C1567R mutant showed enhanced entry into autophagosomes, we examined if its degradation was also increased. Control GECs expressing COL4A5-WT or COL4A5-C1567R were untreated, or incubated with TM, with or without cycloheximide and expression levels were examined after 24 h. Compared to the GECs expressing COL4A5-WT (no TM), the decreases in COL4A5 signals in the presence of cycloheximide were greater in GEC expressing COL4A5-C1567R, and were comparable to COL4A5-WT treated with TM (Fig. 7). Thus, degradation of COL4A5-C1567R was accelerated compared to COL4A5-WT.

**Figure 7.**
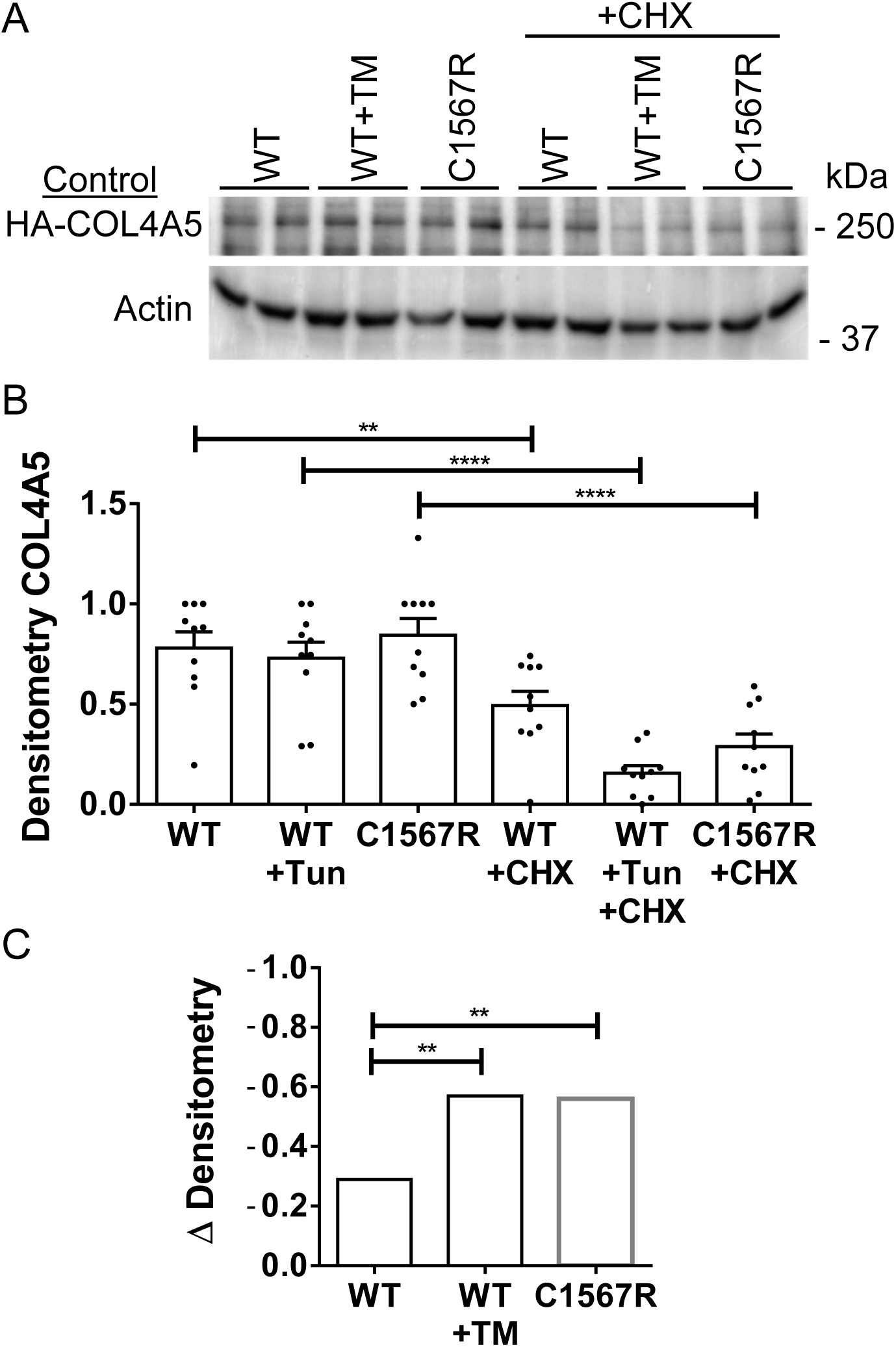
Degradation of the COL4A5-C1567R mutant. Control GECs expressing COL4A5-WT or COL4A5-C1567R were untreated, or incubated with TM (1 μg/mL) with or without cycloheximide (CHX; 1 μM; 24 h). Cell lysates were immunoblotted with antibodies to COL4A5 or actin (A). Signals were quantified by densitometry. B) Densitometric quantification of each group. C) Decreases in densitometric units between the groups of cells treated with cycloheximide and the groups that were not treated. Compared to GECs expressing COL4A5-WT, the decreases in COL4A5 signals in the presence of CHX were greater in GEC expressing COL4A5-C1567R and COL4A5-WT treated with TM. **P<0.01, ****P<0.0001. 6 experiments done in duplicate.

## Discussion

In an earlier study, we had identified Surf4 as a potential non-canonical target of IRE1α [18]. The present study demonstrates that Surf4 can appear in autophagosomes in the context of ER stress. To our knowledge, localization of Surf4 in autophagosomes has not been reported. Basal Surf4 expression was greater in control (IRE1α-replete) GECs compared to IRE1α KO GECs, although Surf4 expression did not increase above basal levels during ER stress. However, glomerular Surf4 protein expression was increased in mice with adriamycin nephrosis in an IRE1α-dependent manner. Furthermore, in human glomerulopathies that involve ER stress and podocyte injury (FSGS and membranous nephropathy), glomerular Surf4 gene expression is increased compared to healthy controls. The finding in cultured GECs did not fully recapitulate the observations in vivo and the reason remains to be elucidated. It should be noted that in a model using inducible Xbp1 in HEK 293 cells, Surf4 protein increased after induction by 1.47-fold above control [44]. Taken together, the IRE1α-Xbp1 axis is involved in Surf4 expression, although some differences may exist between study models.

Further studies in GECs demonstrated that during TM-induced ER stress there was an increase in LC3-II particle area (i.e. autophagosomes) that was partially dependent on IRE1α. This result is in keeping with our earlier studies, which showed that TM increases LC3-II and decreases p62 proteins (by immunoblotting), and increases LC3-II particles (by immunofluorescence microscopy) in an IRE1α-dependent manner [12,13,18]. Surf4 particle area also increased during ER stress, although there was minimal dependence on IRE1α. An increase in colocalization of LC3-II and Surf4 during ER stress implies that a portion of Surf4 was in autophagosomes. Previously, we demonstrated that ER stress increased Sec23B particles, and colocalization of Sec23B with LC3-II [13]. This observation was consistent with the redirection of Sec23B-positive COPII vesicles from the secretory pathway toward autophagy (which was reported to involve phosphorylation of Sec23B by Unc-51 like autophagy activating kinase 1) [21]. By analogy, the present study suggests that a subset of Surf4 was redirected from the secretory pathway toward autophagy.

Podocytes produce structurally-complex proteins that are critical for maintenance of glomerular structure and function [4]. These proteins (e.g. collagens IV) undergo processing in the ER. Our previous study provided evidence that misfolded COL4A5 is degraded by reticulophagy in GECs [13]. Specifically, ER stress (TM) induced the colocalization of COL4A5 and the reticulophagy adaptor RTN3L in autophagosomes. This would potentially reduce accumulation of misfolded collagen IV in the ER. Our model is consistent with findings that COPII vesicles carrying misfolded procollagen I are rerouted from the secretory pathway to autophagosomes and misfolded procollagen I is degraded through reticulophagy [45,46]. The present study further addresses metabolism of misfolded COL4A5 by autophagy. We observed that TM increased COL4A5 and Surf4 colocalization in autophagosomes, suggesting that COL4A5 may be a cargo of Surf4. Indeed, a recent study identified certain fibrillary and non-fibrillar collagens [4] as cargoes of Surf4 in hepatocellular carcinoma cells; collagen IV was not identified, but these cells probably do not express collagen IV [16]. Modeling by Alphafold suggests that Surf4 could potentially act as a receptor or chaperone for the COL4A5 NC1 domain, and that the Surf4 interface spans the three NC1 monomers. This model is novel, and is distinct from another scheme proposed earlier, where Surf4 captures cargo proteins in the ER lumen through an internal pocket, while its cytosolic domain recruits COPII proteins [47]. Our Alphafold model does not take into account that there may be other potential binding partners that optimize the interaction of Surf4 and COL4A5. Thus, the potential interaction of Surf4 and COL4A5 will require further study. During ER stress, COL4A5 was degraded, in keeping with the result of an earlier study [13]. Degradation was attenuated by chloroquine (but not MG132), consistent with a role for autophagy.

Mutations in COL4A3-5 genes cause Alport hereditary nephritis [3,4,48]. Certain collagen IV-α3,4,5 mutant proteins do not form stable trimers, misfold, may be retained in the ER and may activate the UPR in podocytes [36,39]. Such collagen IV mutants cannot assemble a normal GBM. To complement COL4A5 misfolding induced by ER stress, we studied the COL4A5-C1567R mutant. In control GECs expressing COL4A5-C1567R, there were enhanced basal puncta areas of COL4A5, Surf4 and LC3-II, and enhanced COL4A5-C1567R colocalization with Surf4 and with LC3-II compared to basal COL4A5-WT. The COL4A5-C1567R results were similar to the results in COL4A5-WT GECs treated with TM. However, in IRE1α KO GECs, COL4A5-C1567R showed basal puncta areas of COL4A5, Surf4 and LC3-II that were comparable to COL4A5-WT, implying that changes in these parameters in control GECs were at least in part dependent on IRE1α. Colocalization of COL4A5-C1567R with RTN3L in autophagosomes was also greater compared to COL4A5-WT in untreated GECs, and was comparable to COL4A5-WT after induction of ER stress. The results support the view that COL4A5-C1567R is targeted for reticulophagy, and that RTN3L is a reticulophagy adaptor for COL4A5-C1567R (as for endogenous COL4A5-WT) [13]. Finally, degradation of the COL4A5-C1567R mutant in GECs was greater compared to COL4A5-WT, and was comparable to COL4A5-WT in GECs treated with TM. A limitation of the present work is that metabolism of collagen IV in models of glomerular injury in vivo is technically challenging to study. Furthermore, metabolism of collagen IV in diseases is more complex than in cultured GECs and may include isotype switching [4]. Development of additional in vivo models will be required to expand upon results obtained in cell culture systems.

Since podocytes have limited regenerative capacity, preventing podocyte injury is critical to sustaining kidney health. The paucity of effective treatments for glomerulopathies, coupled with evidence that ER stress and impaired proteostasis contribute to their pathogenesis, has recently renewed interest in therapeutically targeting ER stress to attenuate disease progression [49]. In cells, “chemical chaperones” can enhance protein folding and stabilize mutant proteins in the ER, improving export via the secretory pathway and reducing ER stress [5,6,11,50]. We showed this approach can improve protein folding, enhance protein function, and mitigate glomerular injury in experimental GN [51]. In humans, correcting protein misfolding with a chemical chaperone is used as a therapy in Fabry disease [52] and cystic fibrosis [11]. Reducing collagen IV misfolding ameliorated pathology in human kidney organoids [4,53]; however, this approach was not effective in the case of the COL4A5 C1567R mutant [39].

An alternative therapeutic approach could involve stimulation of the UPR or autophagy with drugs. This could alleviate proteotoxic injury by enhancing protein folding capacity and/or degradation of misfolded proteins in podocytes. For example, stimulation of IRE1α RNase with IXA4 in mice was protective in vivo [50,54]. IXA4 enhanced degradation of destabilized amyloid precursor protein variants and reduced cytotoxicity [55]. We showed that in IRE1α-replete GECs, IXA6 (an analog of IXA4) increases expression of ER chaperones [31], thereby increasing ER protein folding capacity. We also identified neratinib as a drug that may stimulate autophagy in GECs [31]. Importantly, we showed that neratinib and IXA6 can reverse adriamycin toxicity in an in vitro assay that reflects podocyte injury in FSGS [31]. A better understanding of the mechanisms of proteostasis in podocytes will allow for further development of therapies that target proteostasis to slow progression of glomerular injury in chronic kidney disease.

## ACKNOWLEDGMENTS

This work was supported by Research Grants from the Canadian Institutes of Health Research (PJ9-166216 and PJ9-169678) and the Kidney Foundation of Canada, and the Catherine McLaughlin Hakim Chair (AVC).

The authors would like to thank Drs. Jeffrey Miner and Kohei Omachi (Washington University, St. Louis, MO) for helpful discussions.

## CONFLICT OF INTEREST

The authors declare that they have no conflict of interest.

## AUTHORS CONTRIBUTIONS

AVC conceived and designed the research, and analyzed data and results. JP, CLKB and RS performed experimental work and examined data and results. AVC wrote the manuscript. All authors reviewed, edited and approved the manuscript.

## Notes

### Competing Interest Statement

The authors have declared no competing interest.

