## Supplemental Figures for "Proteostasis in Podocytes: Role of Cargo Receptor Surf4 and Collagen IV Trafficking"

**Supplemental Figure Legends**

Suppl Figure 1. LC3-II (red) and Surf4 (green) in GECs. See legend to Fig. 3. A) The red and green channels of selected panels from Fig. 3 are presented. B) Changes in puncta numbers and areas induced by TM (with or without chloroquine; CQ) compared to untreated cells. C) In keeping with the particle areas shown in Fig. 3, TM increased LC3-II particle numbers in control GECs, and to a lesser extent in IRE1α KO GECs. **P<0.01, ***P<0.001, ****P<0.0001.

Suppl Figure 2. COL4A5 (red) and Surf4 (green) in GECs. See legend to Fig. 4. The red and green channels of selected panels from Fig. 4 are presented. Arrows point to autophagic puncta.

Suppl Figure 3. Potential interaction of the COL4A5 NC1 domain (amino acids 1457-1685, or the 228 C-terminal amino acids) with Surf4 modeled with Alphafold. The Alphafold structural model (left) predicts NC1 trimer folding and assembly (plDDT) with high confidence, Surf4 folding with moderate to high confidence, and Surf4 docking orientation and binding site(s) with lower confidence. The predicted aligned error plot (right) suggests a protein complex with four regions/chains/domains (A, B, C, D). A, B, and C form three ~228-residue blocks, and is a NC1 domain trimer of COL4A5. A, B, and C have a confidently predicted arrangement. D is Surf4 (~269 residues). AlphaFold is not confident where D sits relative to A, B, and C. Nevertheless, for AlphaFold-Multimer, the ipTM value of 0.68 supports a moderately plausible interface.

Suppl Figure 4. Expression of human COL4A5 mutants. A) Expression of HA-COL4A5-WT and the C1567 and G869R mutants in HEK293T cells. HA-COL4A5-WT and G869R and C1567 mutant cDNAs were expressed in 293T cells by transient transfection. Cell lysates and culture media were immunoblotted with anti-HA or anti-actin antibodies. The two mutants express comparably to COL4A5-WT. COL4A5-WT and G869R are secreted from cells into the culture medium, but C1567R is not. B) 293T cells were untransfected or were transiently transfected with HA-COL4A5-WT and G869R and C1567 mutant cDNAs. After transfection, 293T cells were untreated or treated with TM (0.25 μg/ml; 24 h). The cells were stained with antibodies to HA (COL4A5; red) and calnexin (green). HA staining is evident in transfected cells, but is absent in untransfected cells. C) Colocalization of COL4A5 (red) and LC3-II (green). After transfection, 293T cells were untreated or treated with TM (24 h). Then, cells were stained with antibodies to HA (COL4A5) and LC3. Arrows point to autophagic puncta (yellow), which were observed in WT-expressing cells treated with TM and in cells expressing the mutants. D) Control and IRE1α KO GECs were transduced with lentiviruses to generate clones stably expressing HA-COL4A5-WT. Cell lysates were immunoblotted with antibodies to COL4A5, HA and actin. Untransduced cells, as well as two control and two IRE1α KO transduced clones are presented.

Suppl Figure 5. LC3-II (red) and COL4A5 (green) in GECs. See legend to Fig. 6. The red and green channels of panels from Fig. 6 (left) are presented. Arrows point to autophagic puncta.

Suppl Figure 6. COL4A5 (red) and Surf4 (green) in GECs. See legend to Fig. 6. The red and green channels of panels from Fig. 6 (right) are presented. Arrows point to autophagic puncta.

Suppl Figure 7. Human COL4A5-C1567R mutant in IRE1α KO GECs. IRE1α KO GECs that

,

express HA-COL4A5-WT or HA-COL4A5-C1567R mutant were untreated, or were incubated with TM (0.1-0.25 µg/ml; 24 h). GECs were stained with antibodies to HA (COL4A5; red) and Surf4 (green; left), or with HA (COL4A5; green) and LC3-II (red; right; representative photomicrographs). Puncta area and colocalization of the COL4A5-C1567R mutant with Surf4 and LC3-II were comparable to WT. COL4A5 and Surf4 (left): 4 experiments, N=61-81 per group. COL4A5 and LC3-II (right): 3 experiments, N=75 per group. ****P<0.0001.

Suppl Figure 8. Human COL4A5-C1567R mutant colocalizes with RTN3L. Colocalization of COL4A5-C1567R with RTN3L is greater than COL4A5-WT and is similar to COL4A5-WT treated with TM (control GECs). Puncta area of COL4A5-C1567R tended to increase above COL4A5-WT, but did not reach statistical significance. Control GECs that express HA-COL4A5-WT or HA-COL4A5-C1567R mutant were untreated, or were incubated with TM (0.25 µg/ml; 24 h). GECs were stained with antibodies to HA (COL4A5; red) and RTN3L (green; representative photomicrographs). 3 experiments, N=68-75 per group. *P<0.05, ***P<0.001, ****P<0.0001.

,

Suppl Figure 1

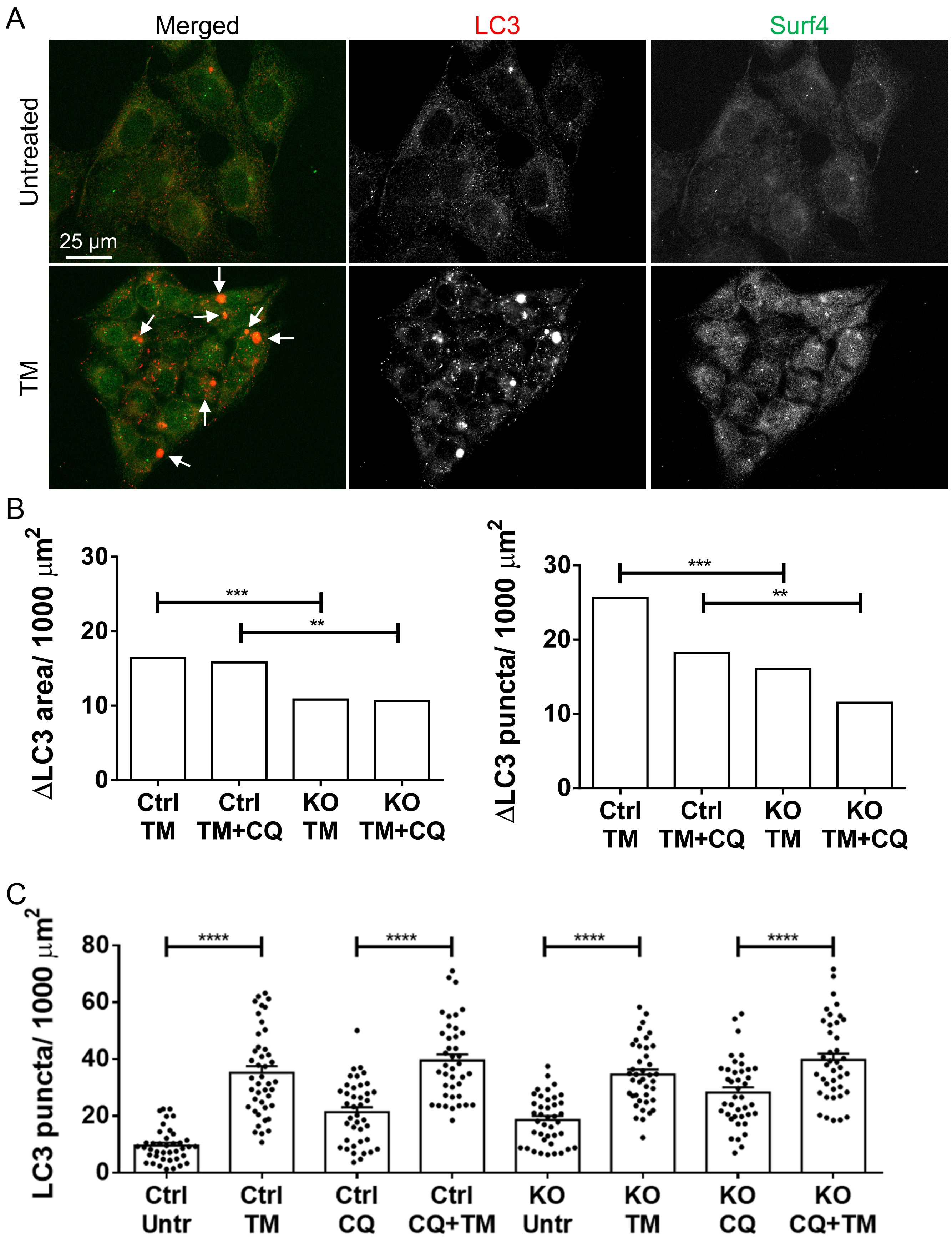

Suppl Figure 2

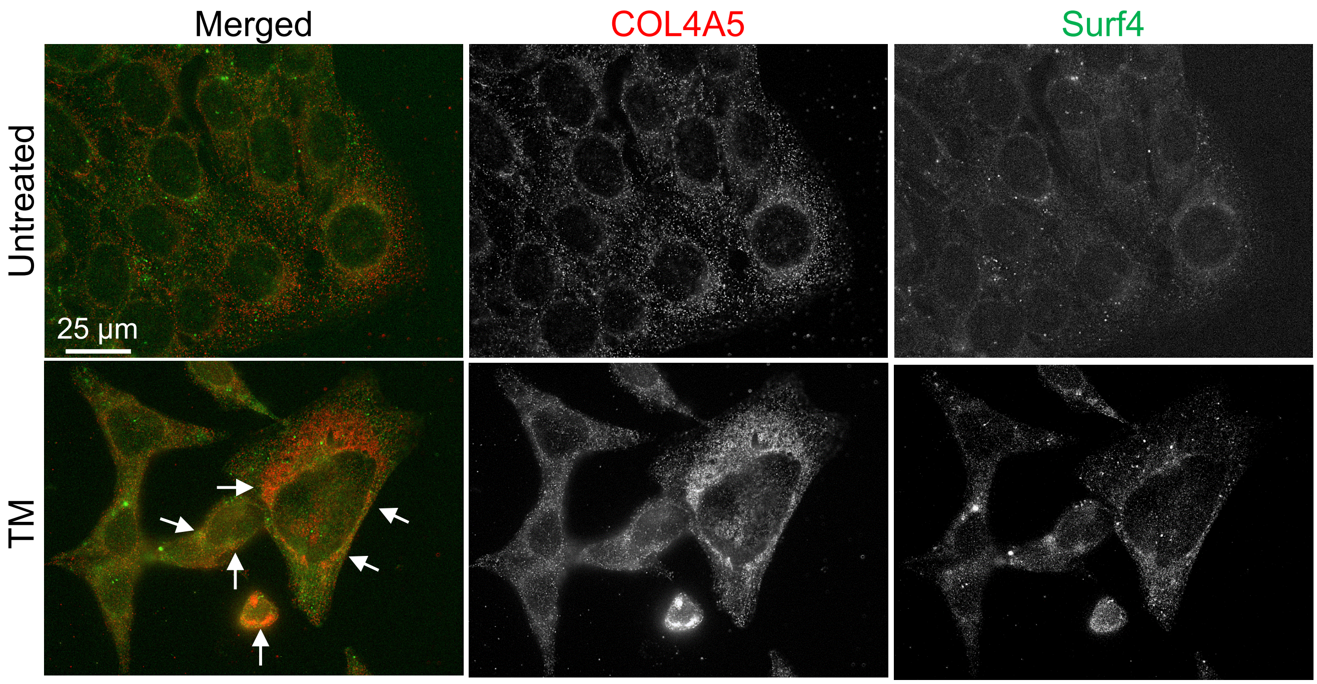

### Suppl Figure 3

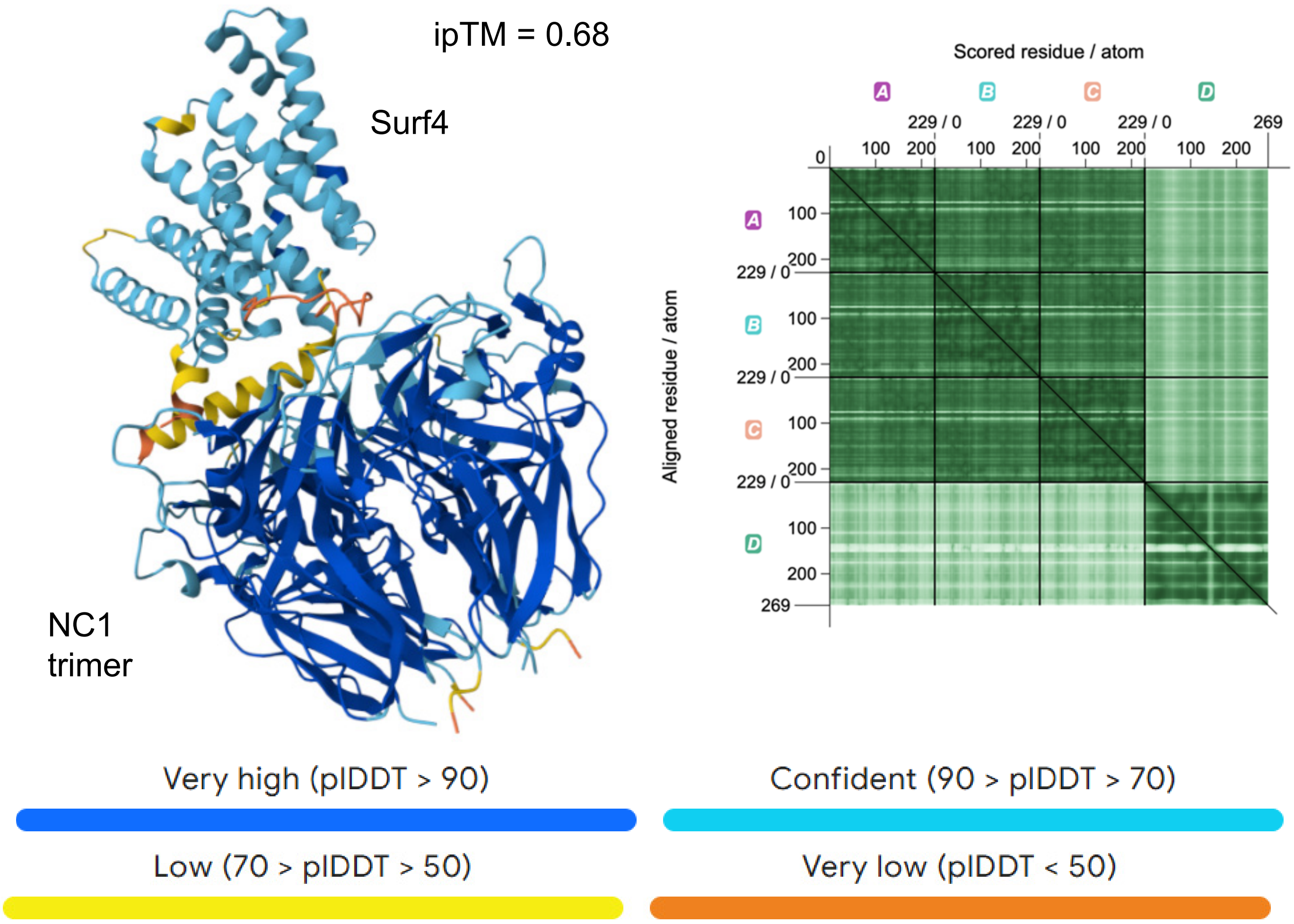

### Suppl Figure 4

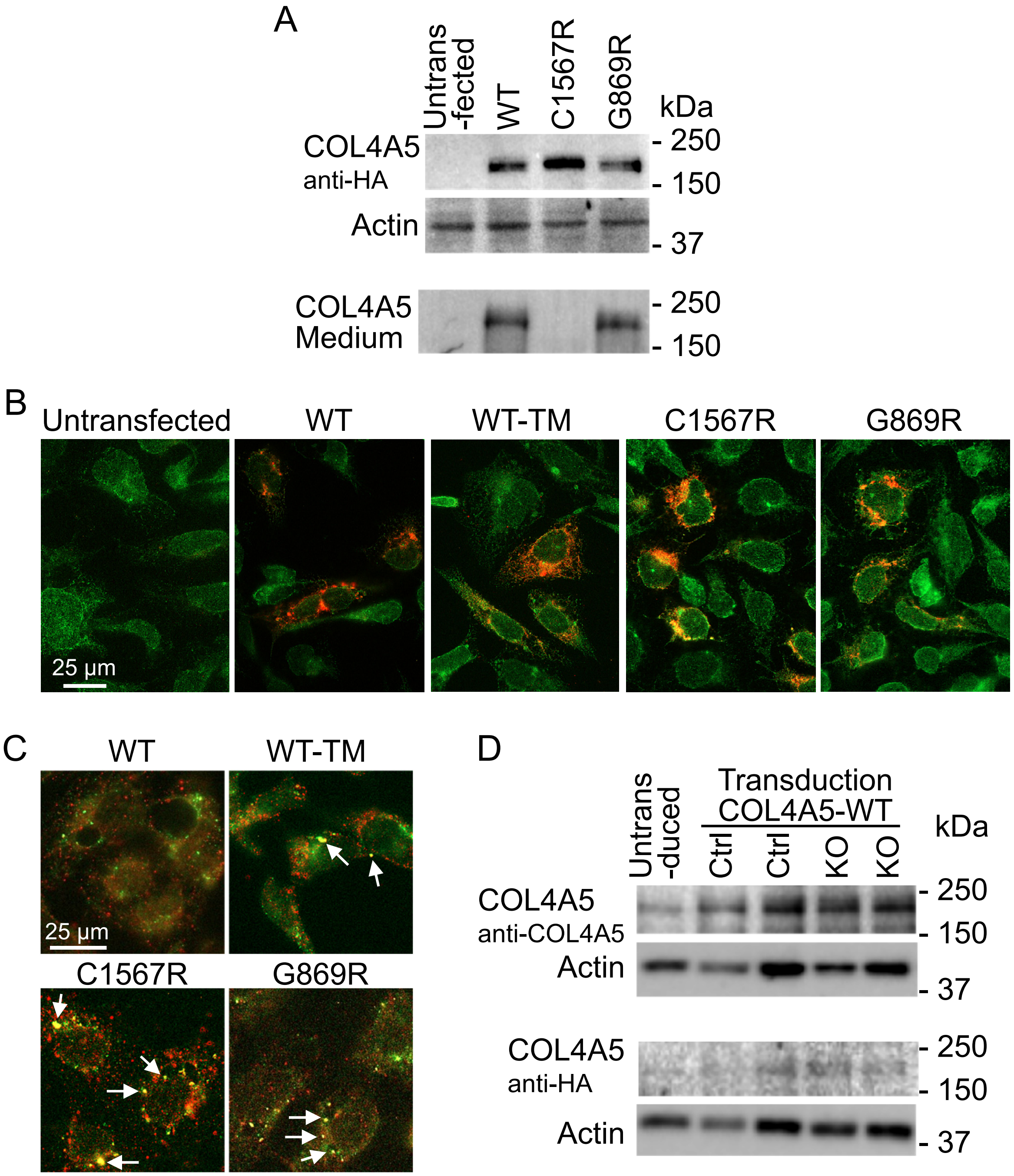

### Suppl Figure 5

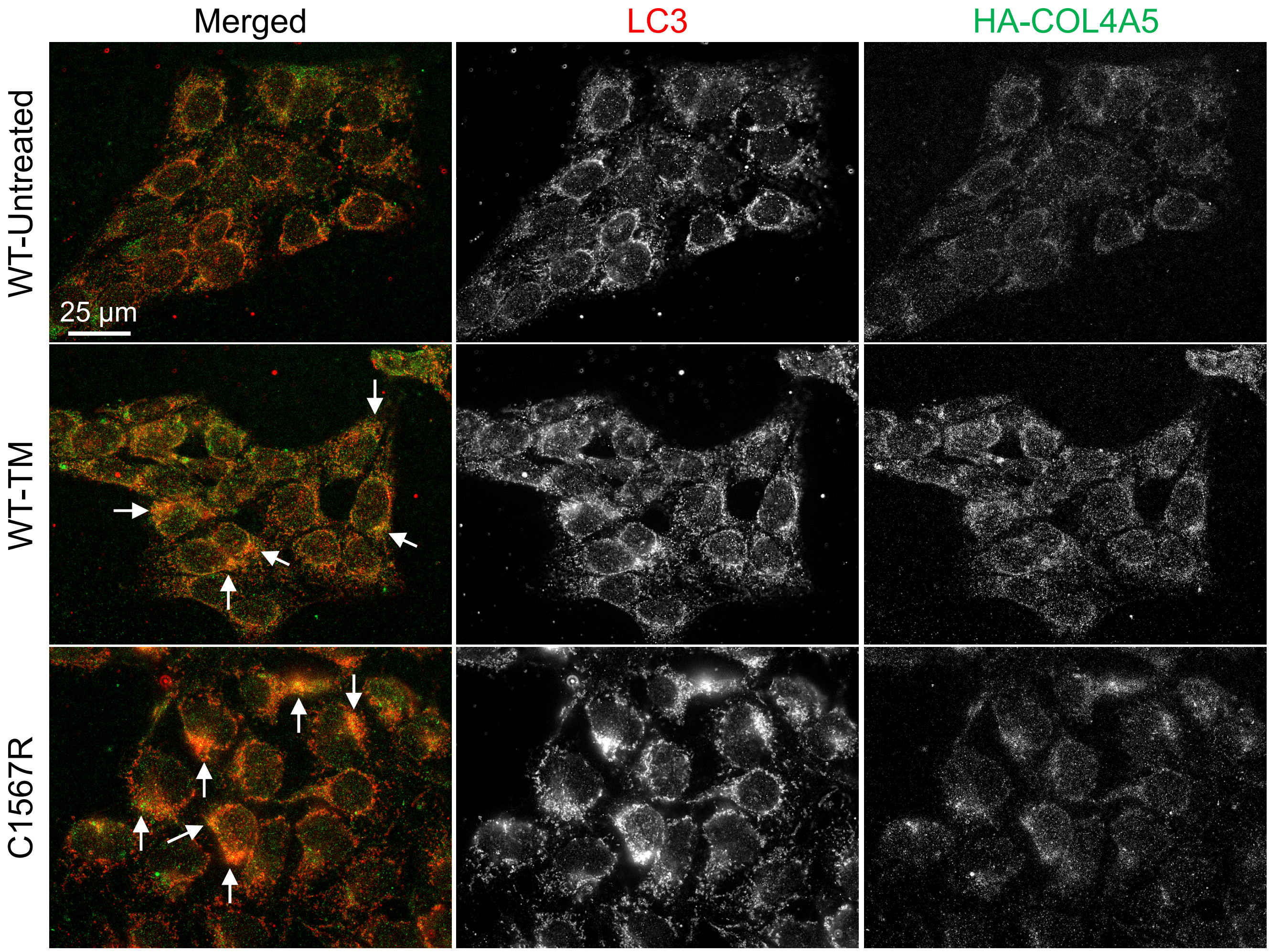

### Suppl Figure 6

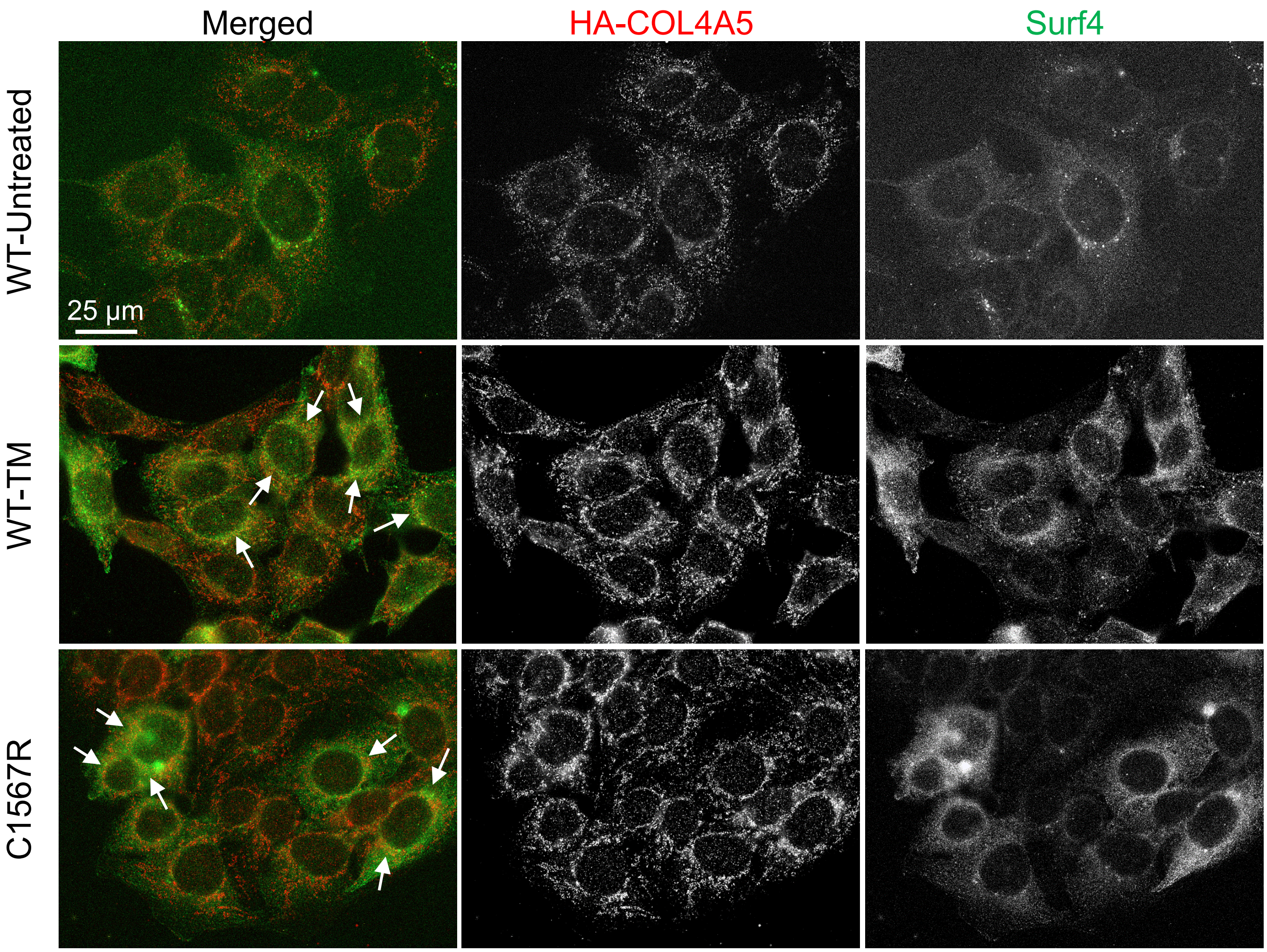

Suppl Figure 7

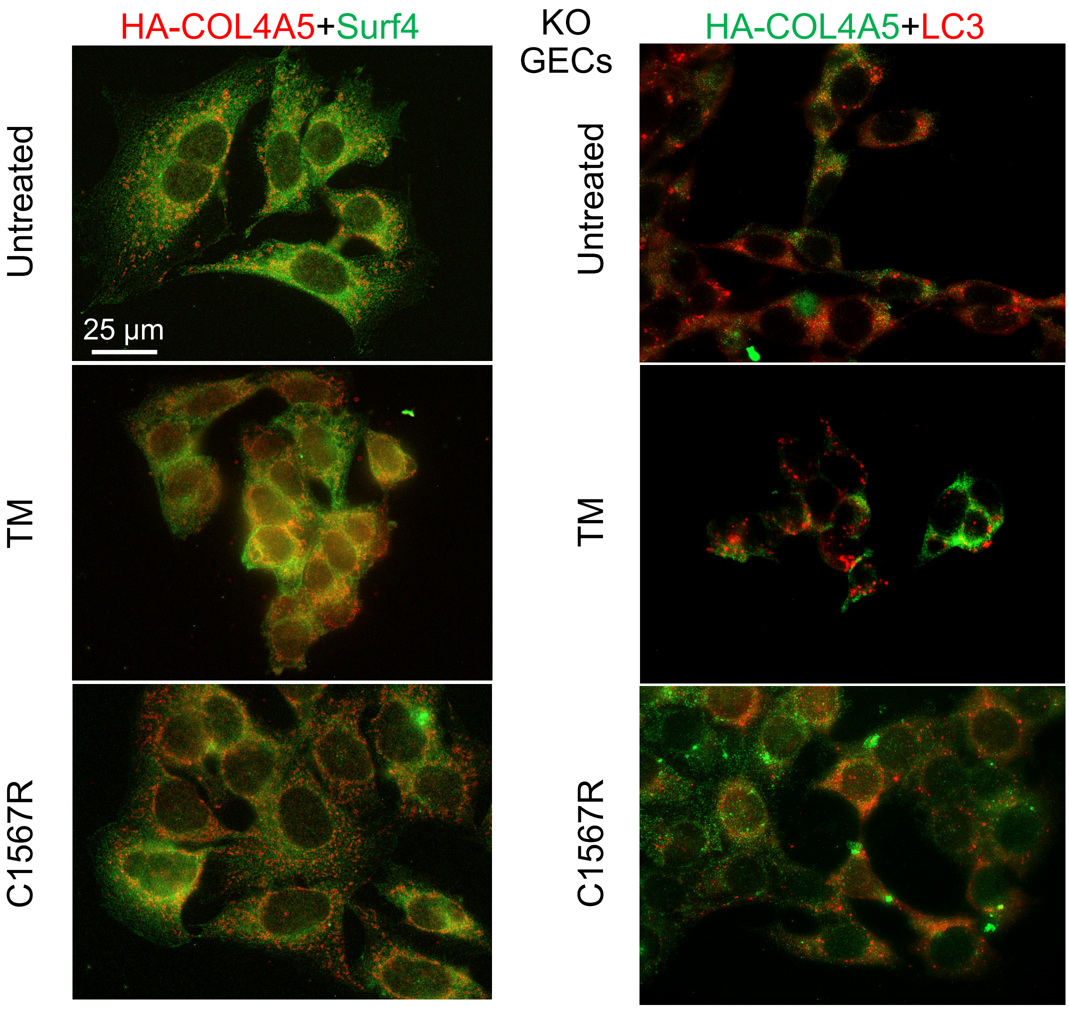

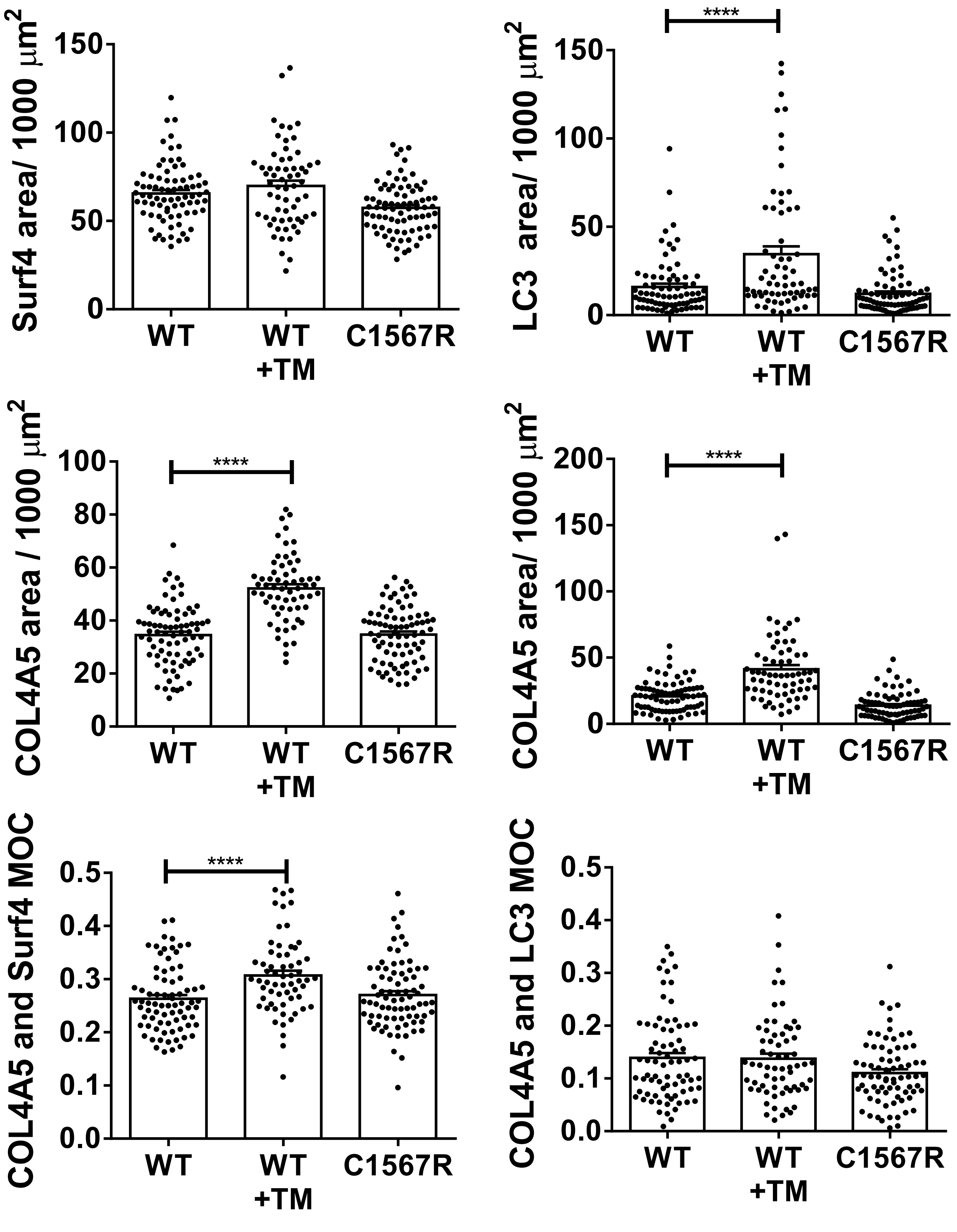
Surf4 area/ 1000 μm2
150
100
50
0
WT
WT +TM
C1567R
LC3 area/ 1000 μm2
****
150
100
50
0
WT
WT +TM
C1567R
COL4A5 area / 1000 μm2
****
100
80
60
40
20
0
WT
WT +TM
C1567R
COL4A5 area/ 1000 μm2
****
200
150
100
50
0
WT
WT +TM
C1567R
COL4A5 and Surf4 MOC
****
0.5
0.4
0.3
0.2
0.1
0.0
WT
WT +TM
C1567R
COL4A5 and LC3 MOC
0.5
0.4
0.3
0.2
0.1
0.0
WT
WT +TM
C1567R

### Suppl Figure 8

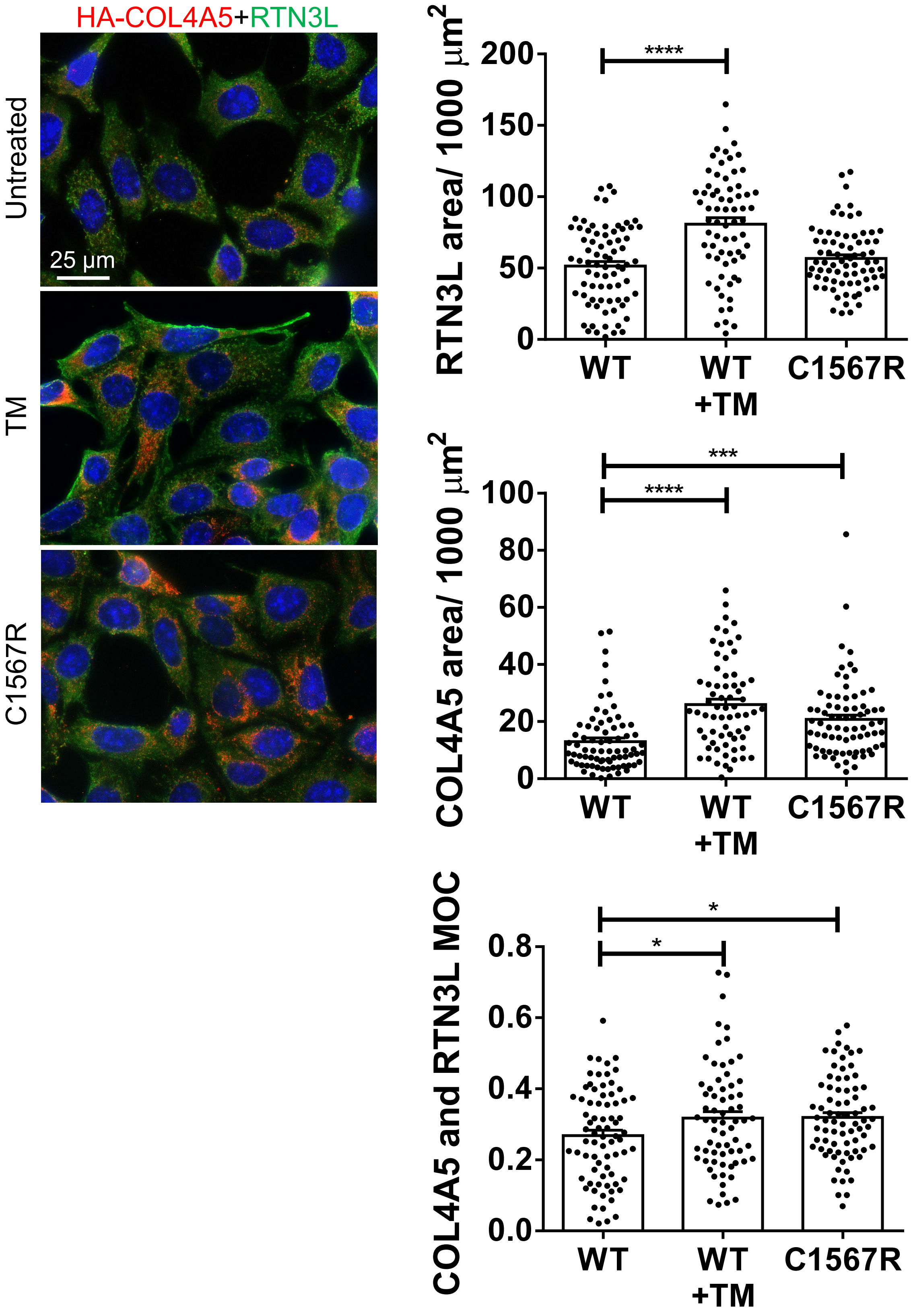
